# HB-EGF Promotes Horizontal Basal Cell Proliferation and Olfactory Sensory Neuron Regeneration in the Zebrafish Olfactory Epithelium

**DOI:** 10.1101/2022.09.26.509477

**Authors:** Siran Sireci, Yigit Kocagöz, Aysu Sevval Alkiraz, Kardelen Güler, Zeynep Dokuzluoglu, Ecem Balcioglu, Mehmet Can Demirler, Stefan Herbert Fuss

**Affiliations:** Bogaziçi University, Department of Molecular Biology and Genetics Center for Life Sciences and Technologies 34342 Bebek / Istanbul, Turkey

**Keywords:** Tissue regeneration, Stem cells, Zebrafish, Olfactory system, Signaling, Neurogenesis

## Abstract

Maintenance and regeneration of the zebrafish olfactory epithelium (OE) are supported by distinct progenitor cell populations that occupy discrete stem cell niches and respond to different tissue conditions. Globose basal cells (GBCs) reside at the inner and peripheral margins of the sensory OE and are constitutively active to replace sporadically dying olfactory sensory neurons (OSNs). In contrast, horizontal basal cells (HBCs) are more uniformly distributed across the tissue, including basal layers of the sensory region, and are selectively activated by acute injury conditions that affect the morphological integrity of the OE. Here we show that expression of the heparin-binding epidermal growth factor-like growth factor (HB-EGF) is strongly and transiently upregulated in response to OE injury and signals through the EGF receptor (EGFR), which is expressed by HBCs. Exogenous stimulation of the OE with recombinant HB-EGF promotes HBC expansion and OSN neurogenesis within the sensory OE, resembling the tissue response to injury. In contrast, pharmacological inhibition of HB-EGF shedding, HB-EGF availability, and EGFR signaling strongly attenuate or delay injury-induced HBC activity and OSN restoration without affecting maintenance neurogenesis by GBCs. Thus, HB-EGF/EGFR signaling appears to be a critical component of a complex signaling network that controls HBC activity and, consequently, repair neurogenesis in the zebrafish OE.

## Introduction

The peripheral olfactory epithelium (OE) is a nervous tissue with an unusually high capacity for cellular self-renewal and repair (**Schwob, 2002**), which makes it an insightful model to study cellular and molecular mechanisms of post-developmental neurogenesis, tissue regeneration, and the regulation of stem cell activity. Neuronal turnover under physiological conditions is sustained by the continuous activity of a dedicated stem/progenitor cell population, the globose basal cells (GBCs; **Calof et al., 2002**; **Krolewski et al., 2013**). In contrast, restoration of the injured OE depends on horizontal basal cells (HBCs), which are largely quiescent in the intact tissue and serve as ‘reserve stem cells’ that are selectively activated by structural damage (**Schnittke et al., 2015**; **Herrick et al., 2017**). In zebrafish, GBCs are restricted to the interlamellar curves (ILCs) and the sensory/non-sensory border (SNS) at the central and peripheral margins of the region that is occupied by olfactory sensory neurons (OSNs), respectively, while HBCs are uniformly distributed across basal layers of nearly the entire tissue (**Bayramli et al., 2017**; **Demirler, 2020**; **Kocagöz et al., 2022**). Accordingly, sustained cell proliferation and OSN generation can be observed at the edges of the sensory OE under physiological conditions (**Bayramli et al., 2017**), whereas the response to injury is characterized by a transient increase in OSN neurogenesis from within the sensory region between the sites of maintenance neurogenesis (**Iqbal and Byrd-Jacobs, 2010**; **Kocagöz et al. 2022**).

A multitude of diverse signaling molecules, including growth factors, neurotransmitters, and small molecules have been described to stimulate or to repress OSN neurogenesis but also to affect HBCs and GBCs differentially (**Buckland and Cunningham, 1998**; **Feron et al., 1999**; **Plendl et al., 1999**; **Simpson et al., 2003**; **Beites et al., 2005**; **Jia et al., 2009**; **Jia and Hegg, 2012**; **Fukuda et al., 2018**). For instance, GBC activity is positively modulated by leukemia inhibitory factor (**Kim et al., 2005**) in the rodent and purine compounds in the zebrafish OE (**Demirler et al., 2020**), whereas growth and differentiation factor 11 (GDF11/BMP11) and Activin βB may serve as negative feedback regulators to ensure optimal OSN density (**Wu et al., 2003**; **Gokoffski et al., 2011**). HBCs in the rodent OE, on the other hand, are activated by changes in Notch/Delta signaling when direct cell-cell contacts between HBCs and sustentacular glial cells (SCs) are disrupted (**Herrick, et al., 2017**). They also respond to tumor growth factor-α (TGF-α), which is expressed by SCs, basal cells, and Bowman’s gland cells but not by OSNs (**Farbman and Buchholz, 1996**). TGF-α is a member of the epidermal growth factor (EGF) family and signals through the EGF receptor (EGFR), which in rodents is expressed by HBCs (**Holbrook et al., 1995**; **Krishna et al., 1996**; **Chen et al., 2020**). It has been shown that EGFR ligands selectively stimulate HBC proliferation but do not affect GBC activity (**Mahanthappa and Schwarting, 1993**; **Farbman and Buchholz, 1996**; **Getchell et al., 2000**). However, evidence for upregulation of EGFR ligands in the injured OE is lacking. More recently, activation of the neural glial-related cell adhesion molecules (NrCAM) in the rodent OE has been implicated in HBC stimulation through an EGFR-dependent mechanism (**Chen et al., 2020**).

Similar to the rodent OE, zebrafish HBCs respond with cell proliferation and OSN neurogenesis when the tissue structure is disrupted but less so when OSNs are selectively ablated by olfactory nerve transection (**Kocagöz et al., 2022**). Activated HBCs give rise to a transient population of mitotically active GBC-like cells, which ultimately restore injured OSNs, eventually other OE cell types. Transcriptome profiling of the regenerating zebrafish OE revealed the prominent activation of diverse signaling pathways, including Wnt/-catenin, cytokine, and TGF-μ signaling, immediately after injury (**Kocagöz et al., 2022**). Among EGFR ligands, a strong and transient upregulation of transcripts for the *heparin-binding epidermal growth factor-like growth factor a* (*hbegfa*; HB-EGF in mammals) and associated matrix-metalloproteases, but not for TGF-α or EGF, could also be observed. Thus, HB-EGF may contribute synergistically or redundantly to HBC activation in the injured zebrafish OE.

HB-EGF, like other EGFR ligands, is activated through ectodomain shedding by matrix metalloproteases (MMPs), most notably by members of the a disintegrin and metalloprotease (ADAM) family (**Higashiyama and Nanba, 2005**; **Blobel, 2005**). The cleaved soluble ectodomain functions as an autocrine and paracrine signal by binding to ErbB1 (= EGFR; **Higashiyama et al., 1991**) and ErbB4 receptors (**Elenius et al., 1997**) to activate a host of intracellular signaling pathways that drive cell proliferation, promote differentiation, and prevent apoptosis (**Singh and Harris, 2005**; **Castanieto et al., 2015**; **Wee and Wang, 2017**). EGFR ligands have been implicated in numerous developmental and regenerative processes by promoting stem cell proliferation and determining lineage identity (**Krampera et al., 2005**; **Dao et al., 2018**; **Abud et al., 2021**). In the nervous system, HB-EGF has been demonstrated to promote neurogenesis during development but also in the postnatal brain in response to various injury conditions (**Oyagi and Hara, 2012**).

Here we show that *hbegfa* is transiently upregulated by OSNs and non-neuronal cells in response to chemical lesion of the zebrafish OE. Changes in *hbegfa* expression are paralleled by increased transcription of *egfra* and members of the ADAM and MMP families that have been demonstrated to contribute to HB-EGF shedding from the membrane. Exogenous stimulation of the OE with recombinant HB-EGF increases mitotic HBC activity and promotes OSN neurogenesis, while inhibition of metalloproteases, soluble HB-EGF, or EGFR signaling severely attenuate or delay injury-induced HBC proliferation and tissue regeneration after injury. Thus, signaling by HB-EGF through EGFR appears to be critically involved in a multifaceted and interdependent signaling network in HBCs that is required for the induction of a full regenerative response in the zebrafish OE.

## Results

### Transcriptional changes in the lesioned zebrafish OE

We have previously described the cellular dynamics and changes in gene expression that occur in the zebrafish OE in response to experimental tissue damage (**Kocagöz et al., 2022**). OSNs undergo rapid degeneration upon nasal irrigation with the detergent Triton X-100 (TrX; **Iqbal and Byrd-Jacobs, 2010**), which reaches a peak at 24 h post lesion (hpl) and is followed by gradual repopulation of the OE with HuC/D-positive sensory neurons over a consecutive 4 to 6 d period. Analysis of RNAseq data (**Kocagöz et al., 2022**; additional data included in this study) revealed that genes typically expressed by OSNs are downregulated over the first 24 hpl but show partial or full recovery by 120 hpl, consistent with the degeneration and reestablishment of OSNs over the same period (**Figure 1A**). Expression of the general neuronal markers *embryonic lethal abnormal visual system-like 3* (*elavl3*; HuC), *microtubule-associated protein 2* (*map2*), and *neuronal cell adhesion molecule 1b* (*ncam1b*) is lowest at 24 hpl but rebounds to pre-injury levels by 120 hpl. Unlike mammals, the zebrafish OE contains multiple morphologically and molecularly distinct OSN subtypes, of which the olfactory receptor/trace amine-associated- (OR/TAAR-) expressing ciliated and the V2R- (= OlfC-) expressing microvillous cells are the most abundant (**Sato et al., 2005**; **Korsching, 2020**). Ciliated OSNs appear to recover more efficiently than apically located microvillous and the fish-specific crypt (*ora4a*), pear (*adora2c*), and *ompa*- or *ora1*-expressing chemosensory cells as indicated by the temporal expression profiles of cell type-specific marker genes (**Sato et al., 2005**; **Oka and Korsching, 2011**; **Oka et al., 2012**; **Suzuki et al., 2015**; **Wakisaka et al., 2017**).

**Figure 1 –.**
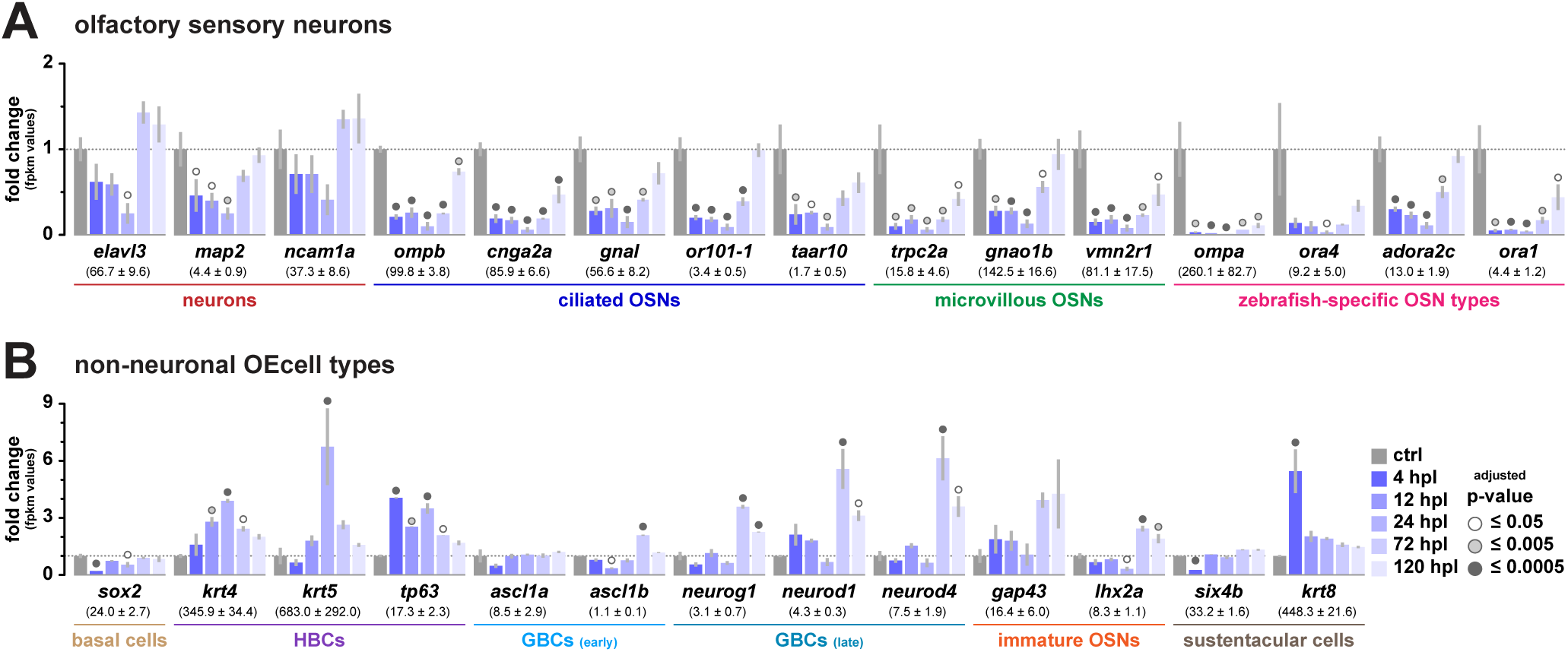
Transcriptional profiling of OE gene expression in response to tissue damage. Changes in gene expression at 4, 12, 24, 72, and 120 h (dark to light blue) following irrigation of the OE with 1% TrX relative to unlesioned control OEs (grey). The data represent whole OE RNA-seq data of three biological replicates for each condition, the numbers below gene names indicate average FPKM values ± SEM of three biological replicates (**Supplementary Table 1**); empty, light, and dark grey circles indicate statistical significance of a Dunnett’s post-hoc test comparing experimental data to controls following a one-way ANOVA at the p ≤ 0.05, p ≤ 0.005, and p ≤ 0.0005 level, respectively. Genome/transcript information, individual FPKM values for three biological replicates, mean FPKM values and statistics information (ANOVA and Dunnett’s *post hoc* tests) are available in Figure 1 – Source data 1. **A.** Transcriptional profiles of genes expressed by olfactory sensory neurons. The *elavl3*, *map2*, and *ncam1a* genes represent general neuronal markers, while ciliated OSNs can be identified by expression of the general markers *olfactory marker protein b* (*ompb*), the *cyclic nucleotide gated channel subunit alpha 2a* (*cnga2*), and the *guanine nucleotide binding protein alpha* (*gnal*), or the subset-specific chemosensory receptors *olfactory receptor 101-1* (*or101-1*) and *trace amine-associated receptor 10* (*taar10*). Microvillous OSN express the *transient receptor potential cation channel subfamily C member 2a* (*trpc2a*) and the *guanine nucleotide binding protein alpha* (*gnao1b*), a subset of which also express the *vomeronasal 2 receptor 1* (*vmn2r1*). The olfactory marker protein a (ompa), *olfactory receptor class A related 4* (*ora4*), *adenosine A2c receptor* (*adora2c*), and *olfactory receptor class A related 1* (*ora1*) genes are specific for unique OSN subtypes of the zebrafish OE. **B.** Transcriptional profiles of cell type-specific non-neuronal markers. In the zebrafish OE, the *SRY box 2* (*sox2*) gene is expressed by basal stem cells (HBCs and GBCs) and sustentacular glial cells. The different non-neuronal population can be individually recognized by the expression of *keratin 4* (*krt4*), keratin *5* (*krt5*), and *tumor protein 63* (t*p63*) in HBCs, *the acheate scute-like* paralogous genes (*ascl1a* and *ascl1b*) by cells of the early GBC lineage, the transcription factors *neurogenin 1* (*neurog1*), *neuronal differentiation 1* (*neurod1*) and *4* (*neurod4*) in late GBCs, *growth-associated protein 43* and *LIM homeobox 2a* (*lhx2a*) in immature OSNs, and the transcription factor *SIX homeobox 4b* (*six4b*) and *keratin 8* (*krt8*) in sustentacular cells. **Figure 1 – Source data 1** **Source data for transcriptional profiling of neuronal and non-neuronal OE cell types** This spreadsheet contains the genome/transcript information, mean FPKM values, individual FPKM values of biological replicates, and statistics information (ANOVA and Dunnett’s *post hoc* tests) for genes shown in Figure 1.

The tissue response to injury is characterized by an early activation of HBCs, which generate a transient population of *ascl1*-positive GBC-like progenitors as regenerative OSN neurogenesis unfolds (**Kocagöz et al., 2022**). At the transcriptional level, the HBC markers *keratin4* (*krt4*) and *krt5* are highly upregulated (3.9 ± 0.1- and 6.7 ± 2.0-fold, respectively) at 24 hpl consistent with HBC activation and increased cell proliferation (**Figure 1B**). The initial response is followed by a modest upregulation of the early GBC markers *ascl1a* and *ascl1b* (1.4 ± 0.2 and 2.1 ± 0.1-fold, respectively) and a strong increase in expression of the late GBC markers *neurogenin 1* (*neurog1*; 3.6 ± 0.1-fold), *neuronal differentiation 1* (*neurod1*; 5.6 ± 1.1-fold), and *neurod4* (6.1 ± 1.2-fold) at 72 hpl. Consistent with the progression of the OSN lineage, the immature OSN markers *growth-associated protein 43* (*gap43*) and *LIM homeobox 2a* (*lhx2a*) are most highly expressed at 72 (*gap43*: 3.9 ± 0.4-fold, *lhx2a*: 2.4 ± 0.1-fold) and 120 hpl (*gap43*: 4.3 ±1.8-fold, lhx2a: 1.9 ± 0.2-fold). In contrast, sustentacular glial cells (SCs) appear to undergo an early reactive gliosis as indicated by the strong increase in expression of the marker *keratin 8* (*krt8;* 5.5 ± 1.1-fold) at the 4 hpl time point despite the small, yet significant, decrease in expression of the alternative zebrafish SC marker *six4b* (**Palominos et al., 2022**, 0.25 ± 0.01-fold), suggesting that some SCs may also undergo degeneration. Thus, the transcriptome profile of OE cell type-specific marker genes closely reflects the events that occur at the cellular level in response to injury (**Kocagöz et al., 2022**), which comprise the rapid degeneration of neurons followed by concomitant activation of HBCs and subsequent GBC activity to generate new OSNs.

### Components of the EGFR/HB-EGF signaling pathway are upregulated in the lesioned OE

We reasoned that relevant signaling pathways, which are involved in the activation of regenerative molecular programs in HBCs, should be upregulated during the early phase of the injury response and precede expansion of the HBC population, which starts between 12 to 24 hpl (**Kocagöz et al., 2022**). Inspection of the transcriptome data revealed the transient upregulation of EGFR signaling components as early as 4 hpl (**Figure 1**). EGFR expression by HBCs has been reported in the rodent OE (**Krishna et al., 1996**; **Ohta and Ichimura, 2000**; **Getchell et al., 2000**; **Gilbert et al., 2015**; **Chen et al., 2020**). EGFR signaling could, thus, be an important target for injury-derived signaling molecules to activate HBC expansion. The zebrafish genome contains six genes that code for EGF receptor subunits, of which *erbb1* (*egfra*), *erbb2*, *erbb3a*, and *erbb3b* show moderate to high expression in the intact tissue, while *erbb4a* and *erbb4b* are only transcribed at low levels (**Figure 2A**). Of those, only *egfra* is selectively upregulated 2.3 ± 0.4-fold at 4 hpl, while expression of the other receptor subunits remains constant (*erbb3b*) or declines during the first 24 hpl to reach pre-injury levels by 72 hpl. The behavior of *erbb2*, *erbb3a*, *erbb4a*, and *erbb4b* subunits, therefore, more closely resembles the temporal changes in gene expression that is characteristic for the various de- and regenerating OSN subtypes (**Figure 1A**). In contrast, *egfra* is more likely transcribed by a cell type that withstands the injury conditions, such as basal progenitor cells (**Kocagöz et al., 2022**).

**Figure 2 –.**
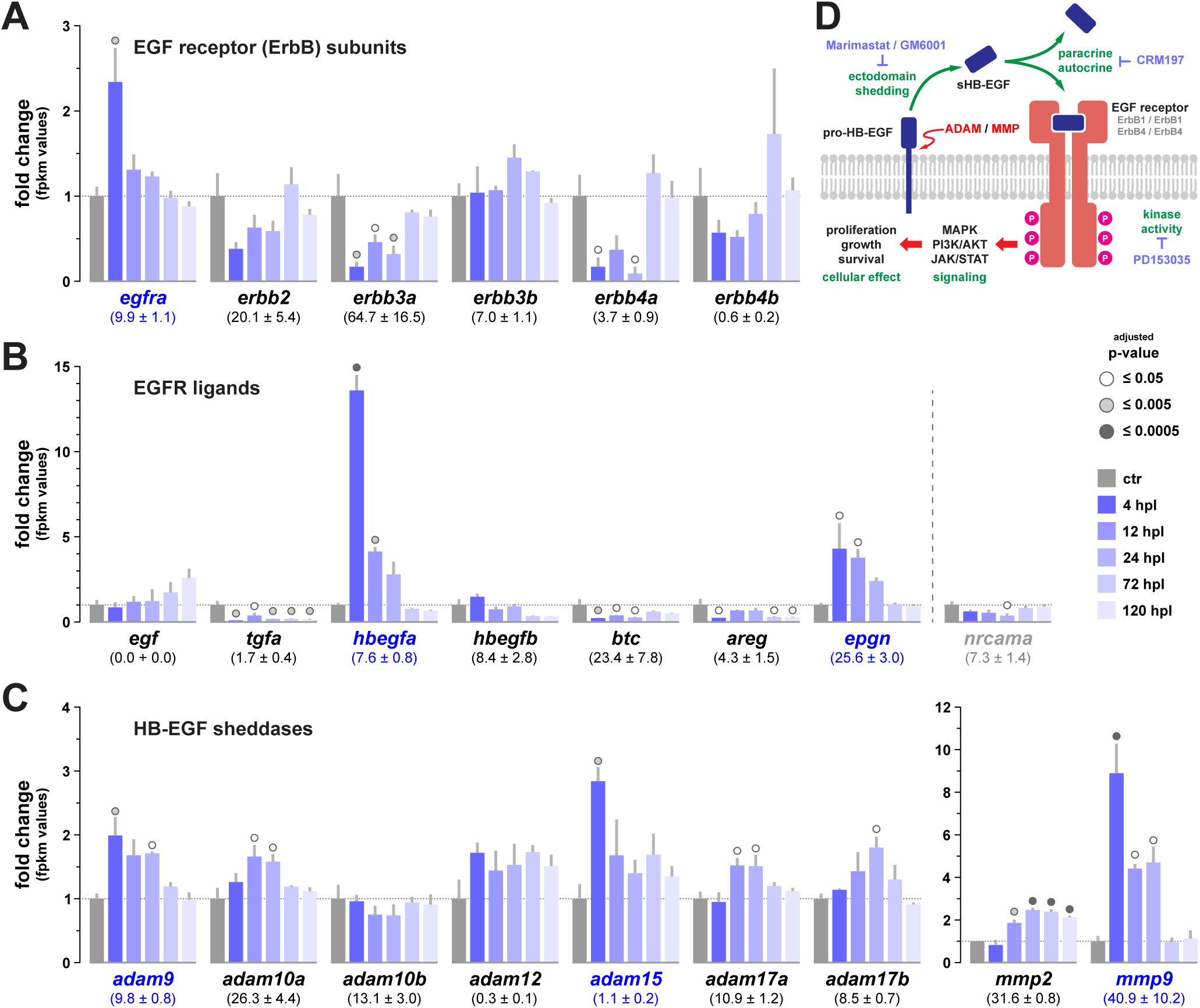
HB-EGF/EGFR signaling is upregulated in response to OE damage. Changes in gene expression at 4, 12, 24, 72, and 120 hpl (dark to light blue) following irrigation of the OE with 1% TrX relative to unlesioned control OEs (grey). The data represent whole OE RNA-seq data of three biological replicates for each condition, the numbers below gene names indicate average FPKM values ± SEM of three biological replicates; empty, light, and dark grey circles indicate statistical significance (adjusted p-value) of Dunnett’s *post-hoc* tests comparing experimental data to controls following one-way ANOVA at the p < 0.05, p < 0.005, and p < 0.0005 level, respectively. Genes that are significantly upregulated at 4 hpl are highlighted in blue. Genome/transcript information, individual FPKM values for three biological replicates, mean FPKM values and statistics information (ANOVA and Dunnett’s *post hoc* tests) are available in **Figure 2 – Source data 1**. **A.** Transcriptional profiles of the six zebrafish ErbB receptor subunits *egfra* (*erbb1*), *erbb2*, *erbb3a*, *erbb3b*, *erbb4a*, and *erbb4b*. *Egfra* is moderately expressed in the intact zebrafish OE and transiently upregulated at 4 hpl. **B.** Transcriptional profiles of EGFR ligands, of which *hbegfa* and *epgn* are transiently upregulated immediately following tissue injury. *Nrcama* is not a canonical EGFR ligand but has been implicated in activating signaling through the receptor. **C.** Transcriptional profiles of matrix metalloproteases that have been demonstrated to function as HB-EGF sheddases, of which *adam9*, *adam15*, and *mmp9* are upregulated early after tissue injury. **D.** Schematic representation of the HB-EGF/EGFR signaling pathway. Pro-HB-EGF is expressed as a single-pass transmembrane protein and activated by cleavage of the extracellular domain by matrix metalloproteases (MMPs) and ADAM sheddases. The soluble ectodomain (sHB-EGF) interacts with ErbB1 or ErbB4 receptors to activate intracellular signaling pathways that stimulate growth, survival, and proliferation. Inhibitors of HB-EGF/EGFR signaling and their molecular targets are indicated by light blue labels. **Figure 2 – Source data 1** **Source data for transcriptional profiling of zebrafish *erbb* genes, ErbB ligands, ADAMs, and metalloproteases** This spreadsheet contains the genome/transcript information, mean FPKM values, individual FPKM values of biological replicates, and statistics information (ANOVA and Dunnett’s *post hoc* tests) for genes shown in Figure 2.

ErbB1 can form homodimeric receptors, which interact with the ligands EGF (*egf*), Tgf-α (*tgfa*), HB-EGF (*hbegfa/b*), betacellulin (*btc*), amphiregulin (*areg*), epiregulin (*epgn*), and epigen (**Linggi and Carpenter, 2006**), which is not annotated in the zebrafish genome. Of those, only *hbegfa* and *epgn* are significantly upregulated 13.6 ± 0.9- and 4.3 ± 1.5-fold at 4 hpl, respectively, while expression of other EGFR ligands remains constant or declines in response to the injury (**Figure 2B**). A recently reported activator of EGF receptor signaling in the mouse OE, the neuron glial-related cell adhesion molecule NrCAM (**Chen et al., 2020**), is not activated following OE lesions in zebrafish and *nrcama* expression declines in a pattern that is similar to genes expressed by OSNs. HB-EGF, epiregulin, and betacellulin can also signal through ErbB4 receptors (**Elenius et al., 1997**; **Fuller et al., 2008**; **Lucas et al., 2022**), and the family of neuregulins are identified ligands for ErbB3 and ErbB4 receptors (**Falls, 2003**). Expression of *neuregulin 1* (*nrg1*) also increases 5.1 ± 2.2-fold at 4 hpl (**Figure 2 – source data 1**). However, ErbB3 and ErbB4 receptors appear not to be relevant in the current context due to their comparably low baseline expression (*erbb3b*, *erbb4a* and *erbb4b*) and/or their strong downregulation at the onset of the injury response (*erbb3a*, *erbb4a*, *erbb4b*). Thus, hbegfa, which is rapidly and strongly upregulated immediately after TrX treatment may constitute a relevant activator of EGFR signaling through Erbb1 homodimers in zebrafish HBCs.

HB-EGF is a single-pass transmembrane protein that is activated by enzymatic cleavage of its soluble extracellular signaling domain, predominantly through members of the ADAM family of metalloproteases (**Figure 2D**). It has been shown that ADAM9, 10, 12, 15, and 17 can function as HB-EGF sheddases in mammals (**Asakura et al., 2002**; **Yan et al., 2002**; **Sahin et al., 2004**) in addition to MMP2, 3, and 9 (**Suzuki et al., 1997**; **Lucchesi et al., 2004**). Of those *adam9*, *adam15*, and *mmp9* are upregulated with a temporal profile that is consistent with their candidate role in HB-EGF activation in the zebrafish OE (**Figure 2C**; see **Figure 2 – Source data 2** for a full complement of zebrafish *adam* and *mmp* gene expression).

### *Egfra* is expressed by HBCs in the zebrafish OE

EGFR is expressed in basal layers in the rodent OE, consistent with its implication in regulating cell proliferation in HBCs (**Krishna et al., 1996**; **Gilbert et al., 2015**; **Chen et al., 2020**). To determine the expression pattern of *egfra* in the zebrafish OE, we performed fluorescent RNA *in situ*-hybridization against *egfra* transcripts in combination with immunohistochemistry against the HBC marker tumor protein 63 (tp63) and the pan-neuronal marker HuC/D (**Figure 3**). Unlike in rodents, the *egfra* riboprobe labeled two distinct cell populations that segregate in basal and apical OE layers, while cells in intermediate strata appear to be devoid of *egfra* expression. The basal *egfra*-expressing cells form a monolayer that is aligned along the lamina propria and co-label with tp63, demonstrating that *egfra* is expressed by HBCs in the zebrafish OE as well. The apical cells, on the other hand, stained positive for HuC/D, implying that *egfra* is also expressed by an uncharacterized OSN subpopulation that is distributed along the apical surface of the tissue. Expression of *egfra* by zebrafish HBCs, however, suggests that EGFR ligands that are upregulated in response to tissue injury may have a direct mitogenic effect on these cells to induce repopulation of the OE with OSNs.

**Figure 3 –.**
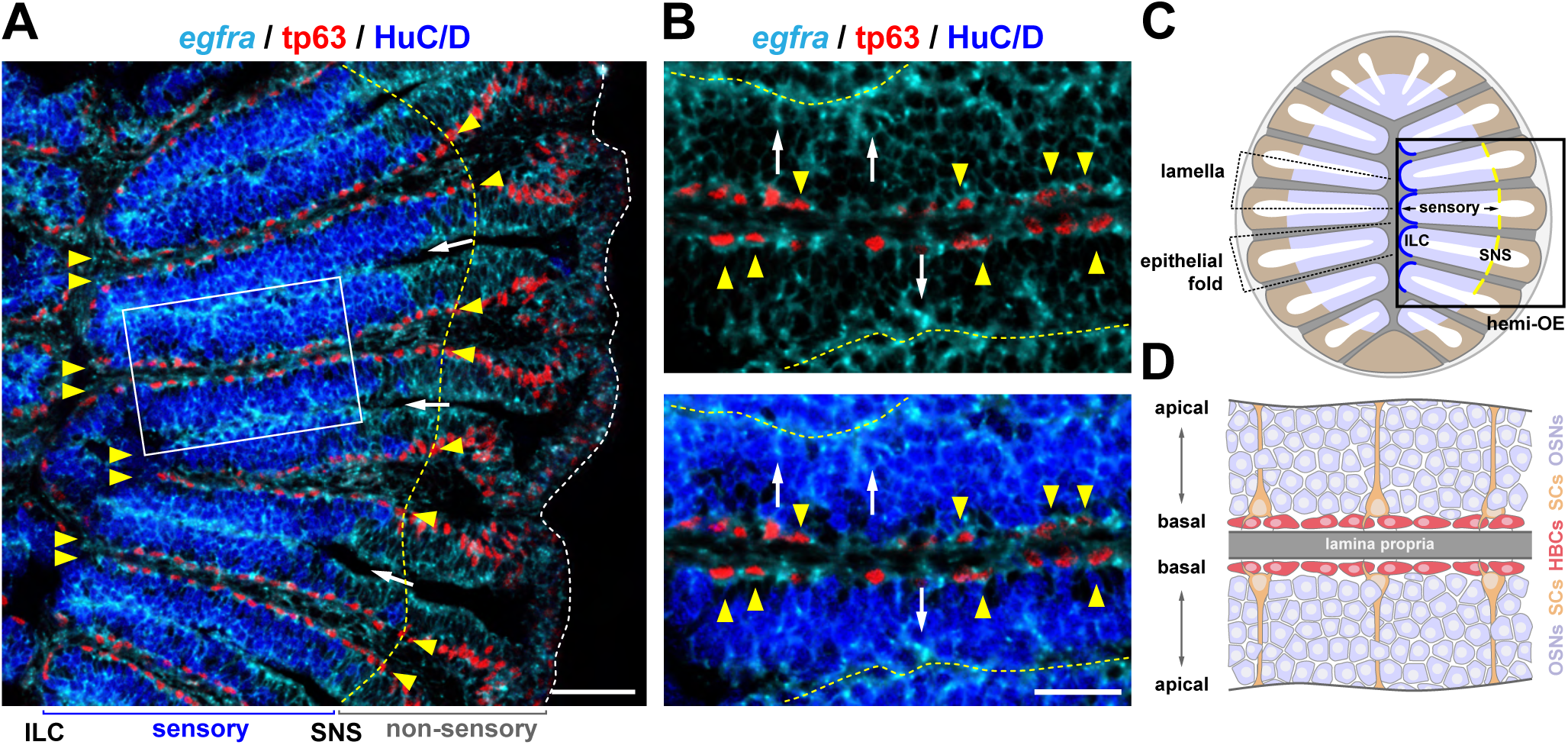
The *egfra* gene is expressed by HBCs in the zebrafish OE. **A.** *In situ*-hybridization against *egfra* transcripts (cyan) and simultaneous detection of the neuronal marker HuC/D (blue) and the HBC marker tp63 (red) by immunohistochemistry on a cross section through the zebrafish OE. Only the right hemi-OE is shown as indicated in C; the interlamellar curves (ILCs) are to the left, the yellow dashed line indicates the sensory/non-sensory border (SNS); the white dashed line demarcates the peripheral outline of the tissue section. Tp63-positive HBCs form double rows in the basal section of each lamella (paired yellow arrowheads) and follow the outline of epithelial folds. The white arrows point towards the water-filled spaces between neighboring lamellae, the light-grey box indicates the region shown in B. Scale bar: 50 µm. **B.** Higher magnification view of the region indicated by the box in A (top: *egfra*/tp63, bottom: *egfra*/tp63/HuC/D). Cells expressing *egfra* segregate into distinct apical and basal layers that stain double-positive for tp63 in the basal OE (yellow arrowheads) and HuC/D in apical strata (white arrows). The yellow dashed lines indicate the apical surfaces of the lamella. Scale bar: 25 µm. **C.** Schematic representation of the zebrafish OE, which is organized into radially projecting lamellae that are formed from two layers of epithelial tissue that are attached to a common lamina propria (grey) at their basal sides. Individual epithelial layers fold back at the ILCs (dark-blue curve) to give rise to U-shaped epithelial folds. Olfactory sensory neurons (OSNs) occupy the inner two-thirds of each fold (blue; sensory) and are surrounded by non-sensory tissue (light brown) with a sharp transition between regions that do and do not contain OSNs at the sensory/non-sensory border (SNS; yellow line). The black box illustrates a hemi-OE similar to the one shown in A and used for quantitative analysis. **D.** Cellular organization of the mid-section of the sensory region of a single lamella as shown in B. Each lamella consists of two epithelial sheets that are connected to a common lamina propria (grey) at their basal sides. Tp63-positive HBCs (red) form a double row of cells along the basal lamina, while sustentacular cell bodies (SC; orange) occupy suprabasal layers and extend processes towards the apical surface. Intermediate and apical layers are predominantly occupied by HuC/D-positive OSNs (blue).

### Expression of the EGFR ligand *hbegfa* is upregulated in the lesioned OE

The transcriptome profile indicates that *hbegfa* is highly and transiently upregulated immediately after OE injury, while the second zebrafish paralog, *hbegfb*, does not show any significant changes in expression (**Figure 2B**). *Hbegfa* transcript levels rise 13.6 ± 0.9-fold by 4 hpl but decline rapidly to 4.1 ± 0.3-fold higher expression by 12 hpl and revert to pre-injury levels between 24 and 72 hpl. However, our RNAseq analysis does not distinguish between different OE cell types. To confirm the RNA sequencing results at tissue level and to gain insight into which cells actively transcribe *hbegfa*, we performed *in situ*-hybridization against *hbegfa* transcripts on intact and lesioned OEs (**Figure 4A**). The *hbegfa* staining pattern in the intact OE was highly variable across animals and tissue sections and only few, if any, *hbegfa*-expressing cells could be detected. If present, *hbegfa*-positive cells almost exclusively occurred as isolated single cells or small cell clusters that were located in close proximity to the basal lamina, thus, the tissue layer where HBCs and SC cell bodies reside (also compare upper left images in panels D, E, and F of **Figure 4**).

**Figure 4 –.**
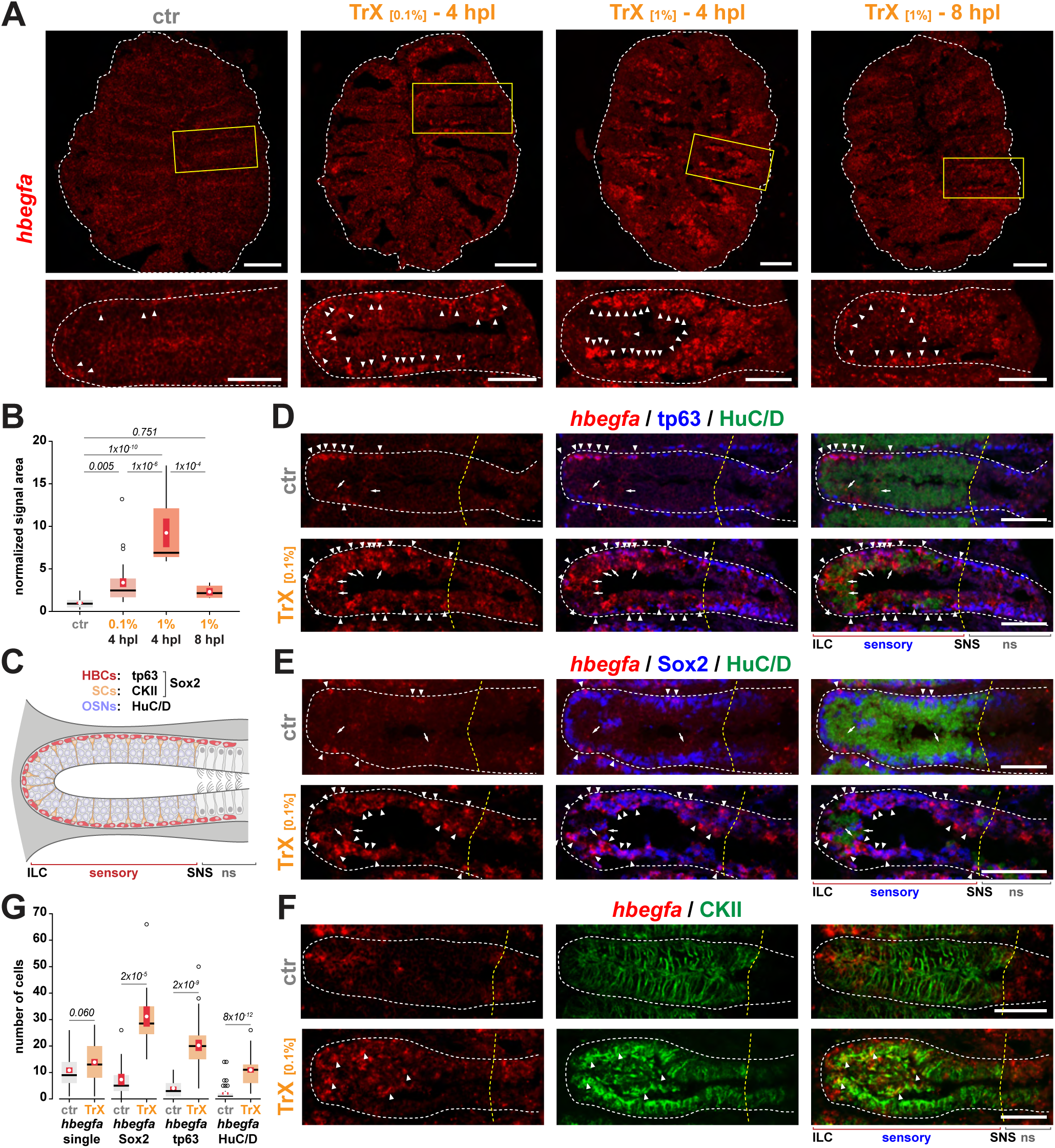
Expression of *hbegfa* is upregulated in the lesioned OE. **A.** *In situ*-hybridization against *hbegfa* transcripts (red) on intact control OEs (ctr, left) and OEs that were lesioned with 0.1% (middle left) or 1% TrX (middle right, right) and analyzed at 4 hpl (left, middle left, middle right), or 8 hpl (right). The top panel shows full OE sections, the outer circumference of the section is depicted by a dashed line. The yellow boxes indicate the regions shown as higher power views in the bottom panel. Individual epithelial folds are outlined by a dashed line; arrowheads depict *hbegfa*-positive cells. Tissue damage induces strong expression of *hbegfa* by 4 hpl, which declines four hours later. Scale bars: 100 µm (top panel), 50 µm (bottom panel). **B.** Quantification of the tissue area covered by *hbegfa*-positive cells. Tissue damage by 0.1% or 1% Trx results in a 3.4 ± 0.5- and 9.2 ± 1.7-fold increase at 4 hpl, respectively. Expression already declines to 2.3 ± 0.4-fold higher levels by 8 hpl in OEs lesioned with 1% TrX. Box and Tukey-style whisker plot; the white dots and red vertical bars represent the means ± SEM of the respective distribution; empty circles denote outliers defined as values ± 1.5 time the interquartile range; p-values indicate the result of Tukey HSD *post-hoc* analysis on one-way ANOVA, F_(3, 59)_ = 23, p = 5.7×10^−10^. The data represent measurements of 24/4 (ctr), 27/3 (4h, 0.1%), 8/1 (4h, 1%), and 4/1 (8h, 1%) hemi-OEs/animals. Raw data for area measurements and statistics information (ANOVA, Tukey HSD post hoc tests) are provided in **Figure 4 – Source data 1**. **C.** Schematic overview of the tissue architecture and distribution of the three major OE cell types that were analyzed in panels D – G; a single epithelial fold is shown. Sox2-positive cells can be subdivided into basal tp63-positive HBCs (red) and suprabasal cytokeratin II-positive sustentacular cells (SCs, orange). OSNs (blue) can be detected by HuC/D expression and occupy the tissue above the SC layer. ILC indicates the interlamellar curve, SNS the sensory/non-sensory border, and ns the non-sensory region that is devoid of OSNs but contains cells that are endowed with beating cilia. **D.** Simultaneous detection of *hbegfa* expression by *in situ*-hybridization (red) and immunohistochemistry against the HBC marker tp63 (blue) and the neuronal marker HuC/D (green) on intact control OEs (ctr, top row) and OEs that have been lesioned with 0.1% TrX (bottom row). The dashed white line demarcates individual epithelial folds as illustrated in C; the yellow line indicates the position of the SNS; arrowheads mark tp63/*hbegfa* double-positive cells, arrows point towards HuC/D/*hbegfa*-positive OSNs. *Hbegfa* is expressed by small clusters of HBCs and occasional OSNs in the intact OE. *Hbegfa* expression expands under injury conditions. Scale bars: 50 µm. **E.** Simultaneous detection of *hbegfa* transcripts (red) and expression of the basal cell marker Sox2 (blue) and the neuronal marker HuC/D (green) by immunohistochemistry on intact control and TrX-lesioned OEs; same design as in D. Arrowheads indicate Sox2/*hbegfa* double-positive cells, arrows demarcate *hbegfa*-positive OSNs. Scale bars: 50 µm. **F.** Simultaneous detection of *hbegfa* transcripts (red) and expression of the SC marker CKII (green) by immunohistochemistry on intact control and TrX-lesioned OEs; same design as in D. The arrowhead indicates a clear double-positive cell. Scale bars: 50 µm. **G.** Quantitative analysis of *hbegf*-expressing cells and overlap with cell type-specific marker expression in intact (ctr, grey) and lesioned (TrX, orange) OEs. Tissue damage induces increased *hbegfa*-expression across all subpopulations. Box and Tukey-style whisker plot; the white dots and red vertical bars represent the means ± SEM of the underlying distribution, respectively; circles denote outliers defined as values ± 1.5 time the interquartile range. The p-values indicate the results of unpaired two-tailed Student’s t-tests for marker-positive cells within individual epithelial folds between control and lesioned OEs (hbegfa single-positive cells: t_(72)_ = −1.9; Sox2: t_(22)_ = −5.43; tp63: t_(48)_ = −7.3; HuC/D: t_(72)_ = −8.14). Cell counts and statistics information (Student’s t-tests) are provided in **Figure 4 – source data 2**. **Figure 4 – Source data 1** **Source data for *hbegfa* area analysis** This spreadsheet provides the raw data for measurements of the area covered by *hbegfa*-expressing cells and statistics information (ANOVA, Tukey HSD post hoc tests) for the data shown in Figure 4B. **Figure 4 – Source data 2** **Source data for quantification of hbegfa-expressing cells and cell type-specific marker coexpression** The spreadsheet contains the cell counts for *hbegfa* single-positive, and Sox2/*hbegfa*, tp63/*hbegfa*, and HuC/D/*hbegfa* double-positive cells alongside statistics information (Student’s t-test) for data shown in Figure 4G.

In contrast, injury by TrX resulted in increased *hbegfa* expression that was both dose- and time-dependent. Mild injury conditions using 0.1% TrX induced a pattern of more uniform expression in basal OE layers at 4 hpl. Both the expression level per cell and the total number of *hbegfa*-positive cells increased further if a higher 1% TrX concentration was used. Under this condition, most of the cells that survived the injury appear to stain positive for *hbegfa*. Consistent with the transcriptome data (**Figure 2B**), *hbegfa* expression already declined by 8 hpl even if a strong 1% TrX stimulus was used. To analyze the effect quantitatively, we thresholded the area occupied by *hbegfa* signal from hemi-OE sections and normalized to the overall size of the tissue (**Figure 4B**). The mild 0.1% TrX application caused a 3.4 ± 0.5-fold increase (p_adj_ = 0.005; Tukey HSD on one-way ANOVA, F_(3, 59)_ = 23.0, p = 5.7×10^−10^), whereas a 9.2 ± 1.7-fold larger area (p_adj_ = 1.4×10^−10^) was occupied by *hbegfa*-positive cells at 4 hpl when the stronger 1% TrX concentration was used. *Hbegfa* expression in 1% TrX-lesioned OEs declined to just 2.3 ± 0.4-fold higher levels by 8 hpl, which was not significantly different from intact control OEs (p_adj_ = 0.751).

Lesions induced by 1% TrX result in severe disruption of tissue morphology, making it difficult, if not impossible, to accurately score individual OE cell types for *hbegfa* expression. To circumvent this problem, we irrigated OEs with 0.1% TrX, which largely preserves the tissue structure but, nevertheless, induces regenerative neurogenesis and HBC activation (**Kocagöz et al., 2022**) and subjected the tissue to simultaneous detection of *hbegfa* transcripts by *in situ*-hybridization and immunohistochemical labeling of the OE with cell type-specific markers at 4 hpl (**Figure 4D, E, F**). As suggested by their basal distribution in the intact tissue, 23.4 ± 3.6% of *hbegfa*-expressing cells also stained positive for the HBC marker tp63 (**Figure 4D, G**), which form a subpopulation of Sox2-positive cells (**Demirler et al., 2020**; **Figure 4E**). The second major Sox2-positive cell population comprises cytokeratin II-positive SCs. However, due to the strong labeling of intracellular filaments, we were not able to score double-positive cells with confidence even in the intact tissue (**Figure 4F**). Nevertheless, the higher number of 36.7 ± 5.1% *hbegfa*/Sox2 double-positive cells suggests that some SCs also express *hbegfa* (**Figure 4G**). In addition, a few *hbegfa*-positive HuC/D double-positive OSNs (between 9.4 ± 1.0%, when scored together with tp63, and 18.3 ± 4.9%, when scored with Sox2) could also be observed.

Tissue damage by 0.1% TrX increased expression of *hbegfa* in all major OE cell types at 4 hpl (**Figure 4G**). The absolute number of *hbegfa* double-positive cells increased 4.3 ± 0.5-fold for Sox2- (ctr: 7.3 ±, 2.1, TrX: 31.2 ± 3.8 cells/epithelial fold; two-tailed unpaired Student’s t-test, t_(22)_ = −5.43, p = 1.9×10^−5^), 5.0 ± 0.5-fold for tp63- (ctr: 4.0 ± 0.6, TrX: 20.2 ± 2.1 cells/epithelial fold; t_(48)_ = −7.31, p = 2.4×10^−9^), and 5.3 ± 0.5-fold for HuC/D-positive cells (ctr: 2.1 ± 0.4, TrX: 10.9 ± 1.0 cells/epithelial fold; t_(72)_ = −8.14, p = 8.3×10^−12^). A reliable quantitative analysis of *hbegfa*/CKII cells was again not possible due to the severe reactive gliosis and altered cell morphology, however, occasional double-positive cells could be identified (**Figure 4F**).

### Exogenous HB-EGF stimulation of the zebrafish OE promotes neurogenic cell proliferation

The intact zebrafish OE is characterized by high proliferative GBC activity at the ILC and SNS, whereas injury also induces neurogenic cell proliferation from within the sensory region of the tissue that is based on HBC activity (**Demirler et al., 2020**; **Kocagöz et al., 2022**). To test whether HB-EGF induces cell proliferation and OSN neurogenesis that resembles injury-induced activity, we stimulated the OE with different concentrations of recombinant human HB-EGF by irrigating one OE with vehicle or recombinant HB-EGF twice for 30 min at 24 h intervals (**Figure 5A**). Following the second HB-EGF application, fish were immediately placed in tank water containing 30 mg/l 5-bromo-2’-deoxyuridine (BrdU) to label mitotic cells for 24 h before analysis by anti-BrdU immunohistochemistry. Irrigation of the OE with vehicle (PBS) or a low 20 ng/µl concentration of HB-EGF did not generate a change in the number or tissue distribution of BrdU-positive cells (**Figure 5B, C**; PBS – ctr: 29.8 ± 2.6, PBS: 27.4 ± 2.6 cell/lamella, two-tailed unpaired Student’s t-test: t_(38)_ = 0.64, p = 0.526; 20 ng – ctr: 47.6 ± 2.2, HB-EGF: 43.2 ± 2.3 cells/lamella, t_(38)_ = 1.38, p = 0.177). However, starting from a concentration of 100 ng/µl HB-EGF, an increased number of BrdU-positive cells could be observed. Stimulation with 100 and 200 ng/µl HB-EGF resulted in a 1.7-fold increase in the overall number of BrdU-positive cells (100 ng - ctrl: 43.3 ± 3.0, HB-EGF: 75.0 ± 5.0 cells/lamella, t_(88)_ = −5.45, p = 4.5×10^−7^; 200 ng – ctr: 18.8 ± 1.1, HB-EGF: 31.2 ± 1.8 cells/lamella, t_(88)_ = −5.83, p = 8.9×10^−8^). The number of mitotically active cells increased evenly across the entire OE and included the occurrence of BrdU-positive cells within the core sensory region (radial positions 0.25 – 0.45; indicated by teal axis labels in **Figure 5B**), in which the number of dividing cells is sparse. Exposure to 100 ng/µl HB-EGF resulted in a 1.8 ± 0.1-fold increase in the number of BrdU-positive cells within the core sensory region (**Figure 5C**; ctr: 8.4 ± 0.9, HB-EGF: 15.3 ± 1.0 cells/lamella, t_(88)_ = −5.10, p = 1.9×10^−6^), while a 2.8 ± 0.4-fold higher number could be recognized upon stimulation with 200 ng/µl of recombinant protein (ctr: 2.1 ± 0.3, HB-EGF: 6.0 ± 0.8 cells/lamella, t_(88)_ = −4.64, p = 1.2×10^−5^).

**Figure 5 –.**
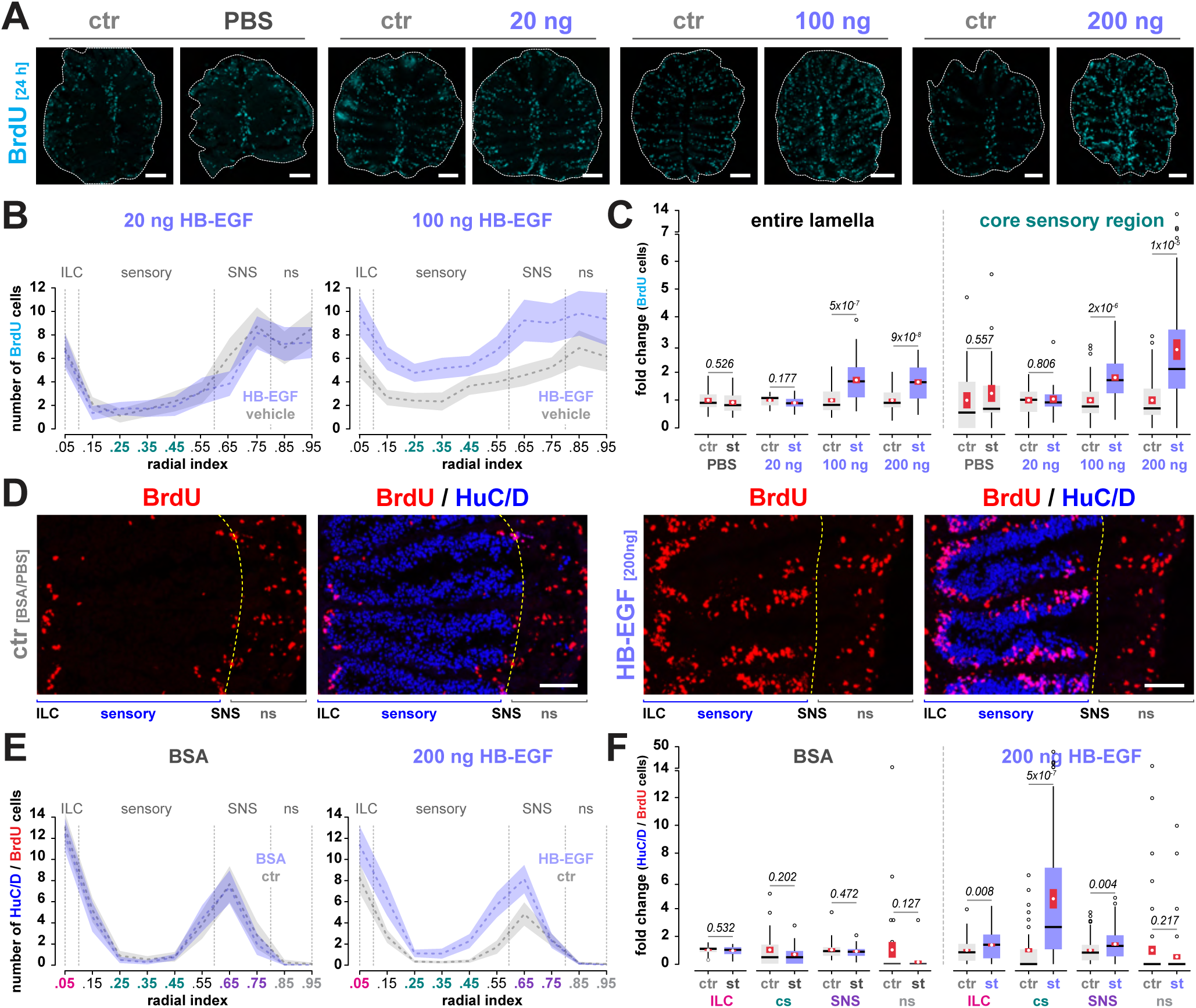
HB-EGF stimulates neurogenic cell divisions in the zebrafish OE. **A.** Immunohistochemistry against BrdU (cyan) on OE sections of fish that were unilaterally treated with vehicle (PBS) or increasing concentrations of HB-EGF (20, 100, 200 ng/µl) and incubated in BrdU-containing water to mark dividing cells for 24 h prior to analysis. The untreated contralateral OEs served as internal control (ctr). The dashed lines mark the outlines of the tissue sections. HB-EGF concentrations higher than 100 ng/µl induce increased cell proliferation in the sensory region of the tissue. Scale bars: 100 µm. **B.** Positional profiling of the distribution of BrdU-positive cells in untreated control OEs and OEs that were stimulated with 20 (left) or 100 ng/µl (right) recombinant HB-EGF. The position of the interlamellar curves (ILCs), the sensory region, the sensory/non-sensory border (SNS), and the non-sensory OE (ns) are indicated; teal x-axis labels mark the core sensory OE analyzed in C. The data represent the mean ± 95% confidence intervals of 20 /5/1 (PBS, 20 ng/µl) or 45/5/3 (100, 200 ng/µl) individual lamellae/OE sections/animals. **C.** Quantification of the effect of HB-EGF stimulation on mitotic activity expressed as the fold change in cell number for individual lamellae between untreated control OEs (ctr) and OEs exposed to different concentrations of HB-EGF (st) over the length of the entire lamella (left) or the core sensory region (right; indicated by teal axis labels in B). Box and Tukey-style whisker plots; the white dots and red vertical bars represent the means ± SEM of the underlying distribution, respectively; circles denote outliers defined as values ± 1.5 time the interquartile range; same data as in B. The p-values denote the results of unpaired two-tailed Student’s t-tests between control and stimulated OEs of the same animals (lamella / core sensory: PBS: t_(38)_ = 0.6 / −0.6, 20 ng: t_(38)_ = 1.38 / −0.3, 100 ng: t_(88)_ = −5.5 / −5.1, 200 ng: t_(88)_ = −5.8 / −4.6). Raw data and statistics information (Student’s t-tests) for B and C are provided in **Figure 5 – Source data 1**. **D.** Immunohistochemistry against BrdU (red) and the neuronal marker HuC/D (blue) on control OEs irrigated twice with the vehicle BSA/PBS (ctr, left) or 200 ng/µl HB-EGF (right). The yellow dashed line indicates the position of the SNS. HB-EGF treatment induces the occurrence of BrdU/HuC/D double-positive cells (magenta) from within the sensory OE. Scale bars: 25 µm. **E.** Positional profiling of HuC/D/BrdU double-positive cells in BSA vehicle-treated control OEs (left, grey) and OEs stimulated with 200 ng/µl HB-EGF (right, light blue). Application of BSA/PBS does not induce OSN neurogenesis, whereas HB-EGF treatment induces neurogenic activity evenly across all OE positions. Colored axis labels indicate the OE regions separately analyzed in F. The data represent the means ± 95% confidence intervals of 30/5/1 (vehicle) and 90/5/3 (HB-EGF) individual lamellae/OEs/animals. **F.** Quantification of the effect of HB-EGF stimulation on OSN neurogenesis expressed as the fold change in cell number between untreated control (ctr) and vehicle- or HB-EGF-stimulated (st) OEs for individual epithelial regions (ILC: interlamellar curve, magenta; cs: core sensory region, teal; SNS: sensory/non-sensory border, purple; ns: non-sensory OE, grey; same data as in E). Application of BSA/PBS did not result in significant changes, whereas stimulation with HB-EGF induced strong neurogenic activity, predominantly within the core sensory region (cs). Box and Tukey-style whisker plot; same data as in E. The p-values denote the results of unpaired two-tailed Student’s t-tests between control and stimulated OEs of the same animals (vehicle – ILC: t_(58)_ = 0.6, cs: t_(58)_ = 1.3, SNS: t_(58)_ = 0.7, ns: t_(88)_ = 1.2; Hb-EGF – ILC: t_(178)_ = −2.7, cs: t_(178)_ = −5.2, SNS: t_(178)_ = −2.9, ns: t_(178)_ = 1.2). Raw data and statistics information (Student’s t-tests) for E and F are provided in **Figure 5 – Source data 2**. **Figure 5 – source data 1** **Source data for HB-EGF concentration effects on cell proliferation** The spreadsheet contains the raw positional cell counts of BrdU-positive cells for vehicle-treated and HB-EGF-stimulated OEs and statistics information (Student’s t-tests) for the data shown in Figure 5B and C. **Figure 5 – source data 2** **Source data for the effect of HB-EGF stimulation on OSN neurogenesis** The spreadsheet includes the raw positional cell counts of BrdU- and BrdU/HuC/D-positive cells for vehicle-treated OEs and OEs stimulated with 200 ng/µl recombinant HB-EGF and staistics information (Student’s t-test) for the data shown in Figure 5E and F.

To examine if the increased mitotic activity is neurogenic and results in the generation of a higher than usual number of OSNs, fish received two consecutive irrigations with 200 ng/µl at a 24 h interval and were immediately placed into BrdU-containing water for 12 h. The animals were then kept for an additional 60 h in clean tank water to allow for some of the marked cells to mature into OSNs before analysis by immunohistochemistry against BrdU and the neuronal marker HuC/D. While vehicle-treated control OEs showed the characteristic pattern of OSN neurogenesis at the ILC and SNS (**Figure 5D**), an increased number of newly born BrdU/HuC/D double-positive OSNs developed from within the sensory region in HB-EGF-stimulated OEs. In total, 1.8 ± 0.1-fold more OSNs were generated in HB-EGF-stimulated OEs (**Figure 5E, F**; ctr: 20.9 ± 1.6, HB-EGF: 38.2 ± 2.5 HuC/D/BrdU-positive cells/lamella, two-tailed unpaired Student’s t-test, t_(178)_ = −5.89, p = 1.8×10^−8^). The effect was again more pronounced in the core sensory region, which usually does not generate a significant number of neurons (**Demirler et al., 2020**) and where HB-EGF stimulation resulted in 4.7 ± 0.7-fold more OSNs (ctrl: 0.9 ± 0.1, HB-EGF: 4.4 ± 0.7 HuC/D/BrdU-positive cells/lamella, t_(178)_ = −5.24, p = 4.6×10^−7^. As seen above for all BrdU-positive cells (**Figure 5B**), the increase in HuC/D/BrdU double-positive cells was again even across all OE regions (**Figure 5E**), suggesting that HB-EGF stimulation activates HBCs, which have a similar tissue distribution, but does not accelerate GBC activity at the ILC and SNS. In contrast, no significant change in the number of HuC/D/BrdU double positive cells was noticeable in vehicle-treated OEs (**Figure 5E, F**; total OE - ctrl: 34.9 ± 2.1, BSA: 32.1 ± 1.8 cells/lamella, t_(58)_ = 1.02, p = 0.312; core sensory - ctr: 2.2 ± 0.5, BSA: 1.5 ± 0.3 cells/lamella, t_(58)_ = 1.02, p = 0.202). Thus, HB-EGF activates a proliferative program that ultimately results in OSN generation in a pattern that resembles regenerative neurogenesis following tissue injury.

### HB-EGF stimulates mitotic activity in HBCs

OSN neurogenesis in the injured sensory OE is powered by increased mitotic activity in HBCs, which gives rise to a transient GBC population that restores the OE (**Kocagöz et al., 2022**). To understand whether exogenous stimulation of the OE with recombinant HB-EGF triggers a similar sequence of events, we stimulated OEs twice with 200 ng/µl HB-EGF, injected fish with 30 µl of 10 mM 5-ethynyl-2′-deoxyuridine (EdU) and stained the tissue samples 24 h after the last HB-EGF stimulation against the proliferation marker and the basal cell marker Sox2 (**Figure 6A**). Sox2 labels a heterogeneous cell population that includes HBCs, SCs, and GBCs. HBCs can be recognized by their horizontal cell profiles that are attached directly to the basal lamina, whereas spherical SCs cell bodies reside in suprabasal layers (**Demirler et al., 2020**). Similar to previous results, HB-EGF irrigation induced a 1.3 ± 0.1-fold increase in overall mitotic activity relative to vehicle-treated control OEs of the same animals (BSA ctr: 18.8 ± 1.3; HB-EGF: 24.2 ± 1.7 EdU-positive cells/epithelial fold; two-tailed unpaired Student’s t-test, t_(88)_ = −2.5, p = 0.015). The increased activity predominantly originated from within the Sox2-positive cell population and included flat-shaped cells that resembled HBCs (**Figure 6A**). Upon HB-EGF stimulation, the number of EdU-labeled Sox2-positive cells more than doubled within the sensory region (**Figure 6B, C**; BSA: 2.6 ± 0.5, HB-EGF: 5.4 ± 0.6 cells/epithelial fold; t_(88)_ = −3.65, p = 4.4×10^−4^), while there was no significant change in the number of single EdU-positive cells that did not stain positive for the Sox2 marker (BSA: 2.7 ± 0.4, HB-EGF: 1.7 ± 0.3 cells/epithelial fold; t_(88)_ = 1.66, p = 0.078).

**Figure 6 –.**
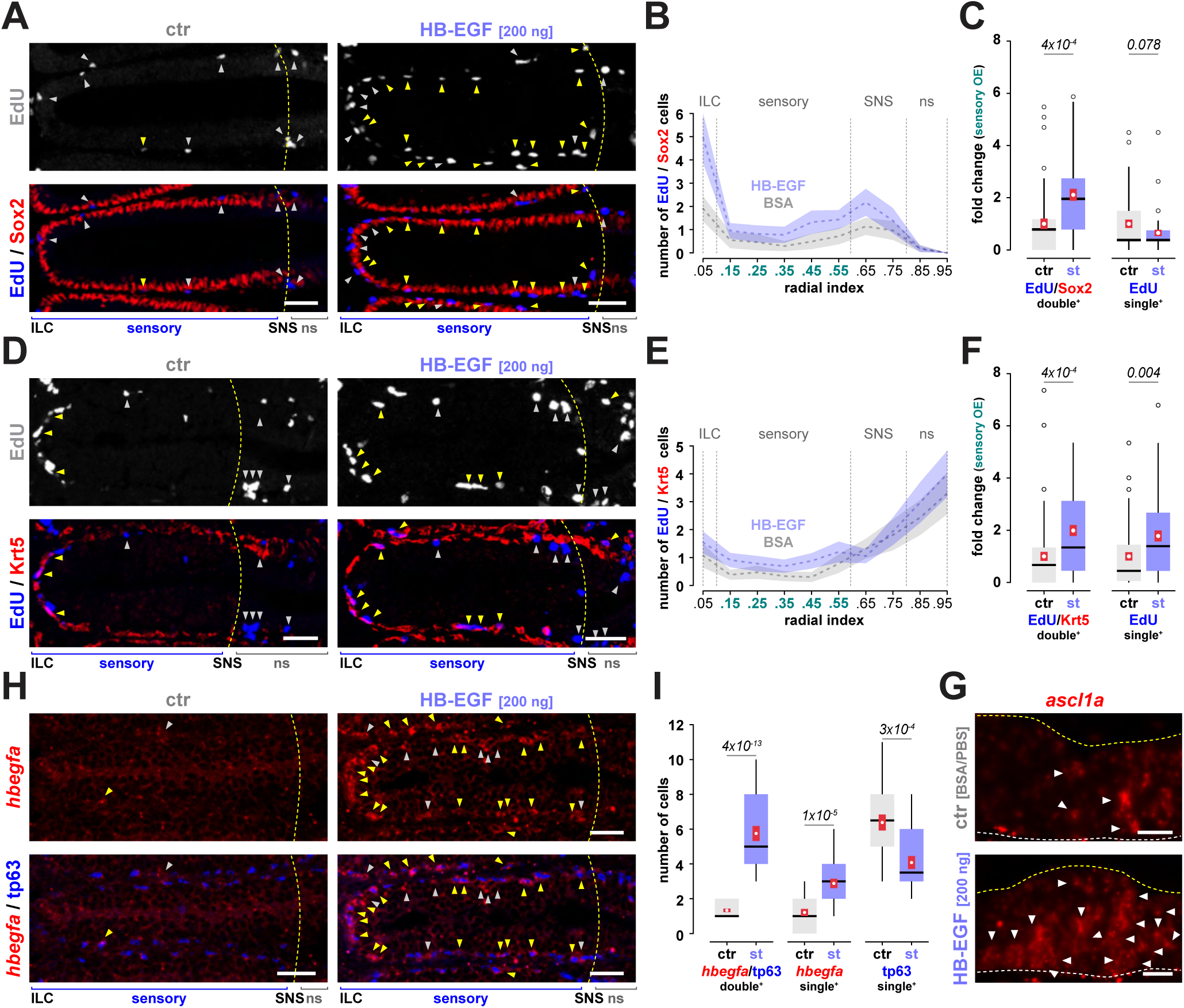
HB-EGF stimulates cell proliferation and *hbegfa* expression in basal cells. **A.** Simultaneous detection of the proliferation marker EdU (white, blue) and the basal cell marker Sox2 (red) on vehicle-treated control OEs (right) and OEs irrigated twice with 200 ng/µl recombinant HB-EGF (right). Fish were injected with EdU immediately after the second HB-EGF stimulation and analyzed 24 h later. HB-EGF induces mitotic activity in the basal OE, including flat-shaped Sox2-positive cells, which resemble HBCs. White arrowheads indicate single EdU-positive cells, yellow arrowheads mark EdU/Sox2 double-positive cells; the yellow dashed line denotes the position of the sensory/non-sensory border (SNS). Scale bars: 25 µm. **B.** Positional profiling of EdU/Sox2 double positive cells in BSA-treated control (grey) and HB-EGF-stimulated OEs (blue). HB-EGF stimulation induces an even increase in the number of mitotically active Sox2-positive cells across all OE positions. The extent of the interlamellar curves (ILC), the sensory region, the sensory/non-sensory border (SNS), and the non-sensory OE (ns) are indicated; teal x-axis labels mark the sensory region analyzed in C. The data represent the means and 95% confidence intervals of 45 individual epithelial folds from 3 fish, 15 folds/animal. **C.** Quantification of the number of EdU/Sox2 double-positive (left) and EdU-single-positive cells in the sensory region of vehicle-treated control OEs (ctr, grey) and OEs irrigated with recombinant HB-EGF (st, blue). HB-EGF preferentially stimulates an increase in the number of mitotically active Sox2-positive cells. Box and Tukey-style whisker plots; the white dots and red vertical bars represent the means ± SEM of the underlying distribution, respectively; circles denote outliers defined as values ± 1.5 time the interquartile range; same data as in B. The p-values denote the results of unpaired two-tailed Student’s t-tests between epithelial folds of stimulated and control OEs (Sox2/EdU: t_(88)_ = −3.7, EdU: t_(88)_ = 1.8). Raw data and statistics information (Student’s t-tests) for B and C are provided in **Figure 6 – Source data 1**. **D.** Immunohistochemistry against EdU (white, blue) and the HBC marker Krt5 (red) on vehicle-treated control OEs (ctr, left) and OEs stimulated with 200 ng/µl HB-EGF; same experimental setup as in A. Yellow arrowheads denote Krt5/EdU double-positive cells, white arrowheads mark single EdU-positive cells; the yellow dashed line denotes the position of the sensory/non-sensory border (SNS). Scale bars: 25 µm. **E.** Positional profiling of EdU/Krt5 double-positive cells in BSA-treated control (grey) and HB-EGF-stimulated OEs (blue). HB-EGF stimulation induces an even increase in the number of mitotically active HBCs across all OE positions. The data represent the means and 95% confidence intervals of 60 individual epithelial folds from 3 fish, 20 folds/animal. **F.** Quantification of the number of EdU/Krt5 double-positive (left) and EdU-single-positive cells in the sensory region of vehicle-treated control OEs (ctr, grey) and OEs irrigated with recombinant HB-EGF (st, blue); same data as in E. HB-EGF preferentially stimulates an increase in the number of mitotically active Krt5-positive and Krt5-negative cells (Krt5/EdU: t_(118)_ = −3.7, EdU: t_(118)_ = −3.0). Raw data and statistics information (Student’s t-tests) for E and F are provided in **Figure 6 – Source data 2**. **G.** *In situ*-hybridization against *ascl1a* transcripts (red) on a vehicle-treated control OE (top) and an OE stimulated once with 200 ng/µl recombinant HB-EGF analyzed at 24 h after stimulation. The mid region of one side of a lamella is shown; the grey dashed line indicates the basal tissue layer, the yellow line the apical surface of the OE. HB-EGF stimulation induces additional *ascl1*-positive cells in the sensory OE. Scale bars: 10 µm. **H.** Detection of *hbegfa* transcripts (red) by RNA *in situ*-hybridization and immunohistochemistry against the HBC marker tp63 (blue) on vehicle-treated control OEs (ctr, left) and OEs irrigated once with 200 ng/µl recombinant HB-EGF (right) 4 h post stimulation. The white arrowheads mark single *hbegfa*-positive cells, yellow arrowheads denote tp63-positive *hbegfa*-expressing cells. HB-EGF induces expression of *hbegfa* in basal cells. Scale bars: 25 µm. **I.** Quantification of hbegfa/tp63 double-positive (left), hbegfa single-positive (center), and tp63 single-positive (right) cells in vehicle-treated control (ctr, grey) and HB-EGF-stimulated (st, blue) OEs. HB-EGF stimulation induces a 3.4-fold induction of *hbegfa* expression in HBCs. Box and Tukey-style whisker plot; the white dots and red vertical lines represent the means and SEMs of the respective distributions from 24 individual epithelial folds of 2 animals, 12 folds/fish. The p-values denote the results of unpaired two-tailed Student’s t-tests for individual epithelial folds between stimulated and unstimulated OEs (*hbefa*/tp63: t_(46)_ = −10.0, *hbegfa*: t_(46)_ = −4.9, tp63: t_(46)_ = 3.9). Raw data and statistics information (Student’s t-tests) are provided in **Figure 6 – Source data 3**. **Figure 6 – Source data 1** **Source data for the quantification of the effect of HB-EGF stimulation on proliferation of Sox2-positive cells** The spreadsheet contains the raw positional cell counts for BrdU-single and BrdU/Sox2 double-positive cells for vehicle-treated and HB-EGF-stimulated OEs and statistics information (Student’s t-tests) for the data shown in Figure 6B and C. **Figure 6 – Source data 2 Source data for the quantification of the effect of HB-EGF stimulation on proliferation of Krt5-positive cells** The spreadsheet contains the raw positional cell counts for BrdU-single and BrdU/Krt5 double-positive cells for vehicle-treated and HB-EGF-stimulated OEs and statistics information (Student’s t-tests) for the data shown in Figure 6E and F. **Figure 6 – Source data 3** **Source data for the effect of HB-EGF stimulation on *hbegfa* expression** The spreadsheet contains the cell counts of tp63 and *hbegfa* single- and double-positive cells for vehicle-treated and HB-EGF-stimulated OEs and statistics information (Student’s t-tests) for the data shown in Figure 6I.

To further discriminate between different cell types, we analyzed the number of Krt5/EdU double-positive cells, which corresponds to the population of dividing HBCs (**Figure 6D**). The number of double-positive cells increased 2.0 ± 0.2-fold across the sensory region in HB-EGF-stimulated OEs (**Figure 6E, F**; BSA: 2.2 ± 0.4, HB-EGF: 4.5 ± 0.5 cells/ epithelial fold; t_(88)_ = −3.65, p = 3.9×10^−4^). Different from the results for Sox2, when scored against Krt5/EdU double-positive cells, the number of single EdU-positive cells also increased 1.8 ± 0.2-fold (BSA: 4.5 ± 0.7, HB-EGF: 8.0 ± 1.0 cells/epithelial fold; t_(88)_ = −2.95, p = 0.004). This observation implies that HB-EGF stimulation of the OE triggers HBC proliferation but also gives rise to additional EdU/Sox2-positive cells, most likely HBC-derived GBCs. To further test this possibility, we performed RNA *in situ*-hybridization against transcripts of the proneural gene *ascl1*, which is expressed by GBCs. Stimulation of the OE with 200 ng/µl HB-EGF also induced *ascl1*-positive cells within the sensory region (**Figure 6G**), which suggests that HB-EGF activates a neurogenic program in HBCs that is identical to injury-induced OSN regeneration.

### HB-EGF stimulates *hbegfa* expression in HBCs

In the zebrafish retina, HB-EGF is part of an autoregulatory gene expression network that includes insulin, growth factors, and cytokines (**Wan et al., 2014**) and it has been suggested that HB-EGF release at the site of injury stimulates *hbegfa* expression in neighboring cells through paracrine signaling routes (**Hashimoto et al., 1994**; **Wan et al., 2012**). Thus, we were curious whether stimulation with recombinant HB-EGF would also affect *hbegfa* expression in the zebrafish OE. Indeed, a single irrigation with HB-EGF induced a robust upregulation of *hbegfa* expression in the basal OE as early as 4 h post stimulation (**Figure 6H**). While in untreated control OEs only 2.5 ± 0.2 *hbegfa*-positive cells could be identified within 80 µm-wide lateral segments of the sensory OE, this number increased more than three-fold to 8.6 ± 0.5 cells (two-tailed unpaired Student’s t-test, t_(46)_ = 12.1, p = 6.8×10^−16^) in HB-EGF-stimulated samples. The majority of cells (66.1 ± 3.0%) with upregulated *hbegfa* expression could be identified as tp63-positive HBCs (**Figure 6I**). The number of *hbegfa*/tp63 double-positive cells increased from 1.3 ± 0.1 to 5.75 ± 0.4 cells/segment (t_(46)_ = −10.0, p = 4.2×10^−13^). Consistent with the increase in double-positive cells, the number of single tp63-positive cells decreased (ctr: 6.4 ± 0.5, HB-EGF: 4.1 ± 0.4 cells/segment; t_(46)_ = 3.9, p = 3.0×10^−4^), while the number of single *hbegfa*-positive cells also increased slightly (ctr: 1.2 ± 0.2, HB-EGF: 2.9 ± 0.3 cells/segment, t_(46)_ = −4.9, p = 1.0×10^−5^). These observations further substantiate that HB-EGF has a direct effect on EGFR-expressing HBCs and results in upregulation of *hbegfa* expression, which then may act through para-/autocrine routes to induce mitotic activity in neighboring HBCs.

### Inhibition of HB-EGF/EGFR signaling prevents injury-induced cell proliferation

HB-EGF signaling is initiated by enzymatic cleavage of its soluble ectodomain, which binds to and activates EGFR (**Higashiyama et al., 2008**). We intercepted this signaling route at three different levels by inhibiting HB-EGF ectodomain shedding, by sequestration of soluble HB-EGF, and by disruption of EGFR activation to test whether HB-EGF signaling is necessary for the induction of mitotic cell divisions in the injured zebrafish OE (**Figure 7**). Cell proliferation and OSN neurogenesis in the intact OE occurs selectively at the ILC and SNS (**Figure 7A** top left; **Bayramli et al., 2017**). As demonstrated before (**Kocagöz et al., 2022**), irrigation of the OE with 1% TrX results in the near-complete loss of HuC/D-positive OSNs by 24 hpl and concomitant activation of cell proliferation across the entire OE, including the occurrence of a high number of mitotically active cells (HBCs and progeny) in the mid sensory region between the ILC and SNS (**Figure 7A** top right).

**Figure 7 –.**
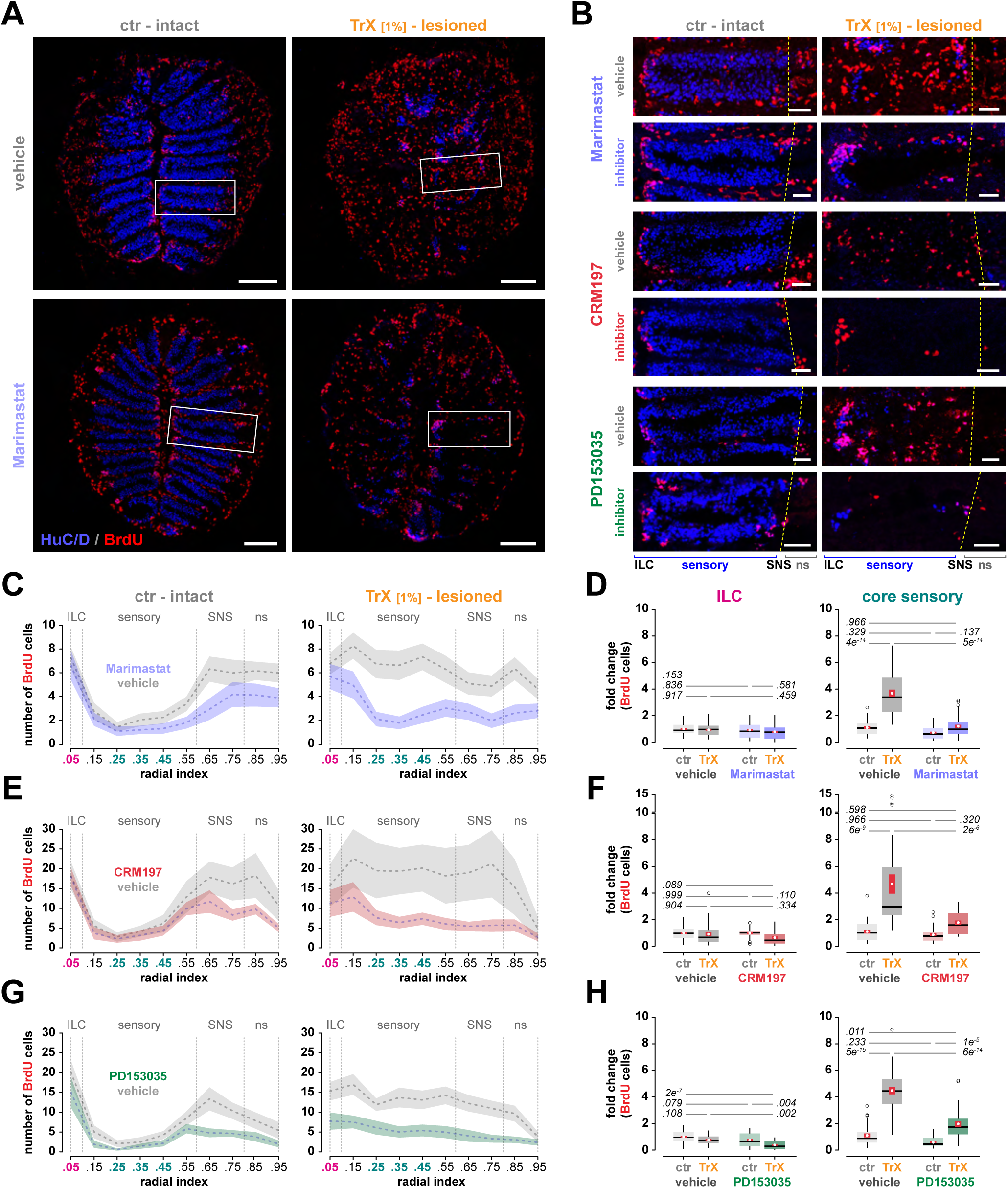
Inhibition of HB-EGF/EGFR signaling prevents injury-induced mitotic activity in the sensory OE. **A.** Immunohistochemistry against BrdU (red) and HuC/D (blue) on intact (left) and TrX-lesioned OEs (right) of fish that were injected with the vehicle DMSO (top) or the broad-spectrum metalloprotease inhibitor Marimastat (bottom). Mitotic activity in the intact OE is high at the interlamellar curves (ILCs) and the peripheral OE flanking the inner sensory region that is occupied by OSNs. TrX treatment results in the destruction of most HuC/D-positive OSNs by 24 hpl and increased mitotic activity in the sensory region of the OE, which is reduced in Marimastat-treated animals. The white boxes indicate the higher power views of individual epithelial folds shown in B. Scale bars: 100 µm. **B.** Immunohistochemistry against BrdU (red) and HuC/D (blue) on intact (left panels) and TrX-lesioned (right panels) OEs of vehicle- and inhibitor-treated fish that were injected with vehicle or the metalloprotease inhibitor Marimastat (blue, top), the HB-EGF inhibitor CRM197 (red, middle), and the EGFR inhibitor PD153035 (green, bottom). Individual epithelial folds are shown, the ILCs are to the left, the non-sensory OE (ns) to the right. The dashed yellow line demarcates the position of the sensory/non-sensory border (SNS). All three inhibitors prevent the unfolding of injury-induced mitotic activity in the sensory region of the OE. Scale bars: 25 µm. **C.** Positional profiling of BrdU-positive mitotic cells in intact control OEs (left) and TrX-lesioned OEs (right) of fish that were injected with vehicle (grey) or Marimastat (blue), which prevents induction of injury-induced mitotic activity in the lesioned OE. Colored axis labels indicate the OE regions separately analyzed in D.The data represent the means and 95% confidence intervals of 45 individual lamellae from 3 animals, 15 /lamellae/fish for each condition. **D.** Quantification of the number of mitotically active cells in the ILC (left; indicated by magenta axis labels in C) and the core sensory OE (right; teal) in untreated control (ctr) and TrX-lesioned OEs (TrX) of fish that were treated with vehicle (grey boxes) or Marimastat (blue boxes) expressed as the fold change in the number of BrdU-positive cells relative to intact control OEs of vehicle-treated animals. Marimastat treatment does not significantly affect ongoing cell proliferation in the ILC but severely diminishes mitotic activity in the core sensory region of lesioned tissues. Box and Tukey-style whisker plot; the white dots and red vertical lines represent the means and SEMs of the respective distributions; same data as in C. The p-values denote the results of Tukey HSD *post-hoc* for meaningful comparisons following one-way ANOVA (ILC: F_(3, 176)_ = 1.6, p = 0.199; core sensory: F_(3,176)_ = 84.6, p = 2×10^−16^). **E.** Positional profiling of BrdU-positive mitotic cells in intact and TrX-lesioned OEs of vehicle-injected control fish (grey) and animals in which soluble HB-EGF was sequestered by CRM197 (red). Inhibition of HB-EGF signaling prevents induction of injury-induced mitotic activity in the lesioned OE. The data represent the means and 95% confidence intervals of BrdU-positive cells of 30 individual hemi-OEs from 3 animals, 10 hemi OEs/fish for each condition. **F.** Quantification of the number of mitotically active cells in the ILC (left) and the core sensory OE (right) in intact (ctr) and TrX-lesioned OEs (TrX) of fish that were treated with vehicle (grey boxes) or CRM197 (red boxes); same data as in E; design as in D. CRM197 treatment does not significantly affect ongoing cell proliferation in the ILC but severely diminishes mitotic activity in the core sensory region. The p-values denote the results of Tukey HSD *post-hoc* tests on one-way ANOVA for meaningful comparisons following one-way ANOVA (ILC: F_(3, 116)_ = 2.4, p = 0.071; core sensory: F(3,176) = 21.6, p = 3.6×10^−11^). **G.** Positional profiling of BrdU-positive mitotic cells in intact control and TrX-lesioned OEs of vehicle-injected fish (grey) and animals in which EGFR signaling was inhibited by PD153035 (green). PD153035 diminishes induction of injury-induced mitotic activity in the lesioned OE. The data represent the mean and 95% confidence intervals of BrdU-positive of 30 individual hemi OEs from 3 animals, 10 hemi-OEs/fish for each condition. **H.** Quantification of the number of mitotically active cells in the ILC and the core sensory OE in untreated control (ctr) and TrX-lesioned OEs (TrX) of fish that were treated with vehicle (grey boxes) or PD153035 (green boxes); same data as in G; design as in D. PD153035 also diminishes mitotic activity in the core sensory region. The p-values denote the results of Tukey HSD *post-hoc* tests for meaningful comparisons following one-way ANOVA (ILC: F_(3, 116)_ = 11.9, p = 7.3×10^−7^; core sensory: F_(3,116)_ = 78.9, p = 2×10^−16^). **Figure 7 – Source data 1** **Source data for the effect of inhibition of metalloprotease activity by Marimastat on injury-induced cell proliferation at 24 hpl** The spreadsheet contains the raw positional cell counts for BrdU-positive cells in intact control and 1% TrX-lesioned OEs of vehicle- and Marimastat-treated animals alongside statistics information (one-way ANOVA, Tukey HSD *post hoc* tests) for the data shown in Figure 7C and D. **Figure 7 – Source data 2** **Source data for the effect of HB-EGF sequestration by CRM197 on injury-induced cell proliferation at 24 hpl** The spreadsheet contains the raw positional cell counts for BrdU-positive cells in intact control and 1% TrX-lesioned OEs of vehicle- and CRM197-treated animals alongside statistics information (one-way ANOVA, Tukey HSD *post hoc* tests) for the data shown in Figure 7E and F. **Figure 7 – Source data 3** **Source data for the effect of inhibition of EGFR activity by PD153035 on injury-induced cell proliferation at 24 hpl** The spreadsheet contains the raw positional cell counts for BrdU-positive cells in intact control and 1% TrX-lesioned OEs of vehicle- and PD153035-treated animals alongside statistics information (one-way ANOVA, Tukey HSD *post hoc* tests) for the data shown in Figure 7G and H.

To prevent HB-EGF ectodomain shedding, we utilized the broad-spectrum MMP inhibitor Marimastat (**Rasmussen and McCann, 1997**; **Figure 7A, B** top panel). Fish were primed by two intraperitoneal (IP) injections of 80 µg/g Marimastat or DMSO vehicle at 24 and 12 h prior to lesioning the tissue with 1% TrX and received additional inhibitor injections at the time of TrX treatment and at 12 hpl. To label mitotic cells, animals were placed in BrdU-containing water immediately after the lesion until analysis at 24 hpl. Tissue damage by TrX treatment in vehicle-injected control fish disrupted the bimodal distribution of dividing cells and induced an overall 1.5 ± 0.1-fold increase in the number of BrdU-positive cells (**Figure 7C**; vehicle - ctr: 43.5 ± 1.95, TrX: 63.0 ± 3.2 cells/lamella; p_adj_ = 1.6×10^−6^, Tukey HSD *post-hoc* analysis on one-way ANOVA, F_(3,176)_ = 37.6, p < 2×10^−16^). In the core sensory region, a 3.6 ± 0.3-fold higher number of BrdU-labeled cells could be observed (**Figure 7D**; vehicle - ctr: 5.7 ± 0.5, TrX: 20.7 ± 1.5 cells/lamella; p_adj_ = 4.1×10^−14^, F_(3,176)_ = 75.5, p < 2×10^−16^). Treatment with Marimastat had a pronounced effect on injury-induced cell proliferation. Overall, the BrdU staining pattern in the lesioned OE of inhibitor-treated fish more closely resembled the mitotic activity pattern in the intact tissue with residual cell proliferation at the ILC and SNS but only sporadic activity in the sensory region (**Figure 7A, C**). In the presence of Marimastat only a non-significant 1.1 ± 0.1-fold increase in the number of BrdU-positive cells could be observed in the core sensory region relative to the intact OE of DMSO-injected control fish (Marimastat – TrX: 6.3 ± 0.7 cells/lamella; p_adj_ = 0.966; **Figure 7C, D**). Marimastat treatment had no obvious effect on maintenance neurogenesis at the ILC, either in the intact or lesioned tissue at the observation time point (**Figure 7D**). Thus, inhibition of MMP activity prevents the typical early response to tissue damage, which goes along with a selective increase in HBC cell proliferation in the sensory region. However, it should be noted that Marimastat has a broad target spectrum and does not selectively inhibit MMPs that act on HB-EGF but also inhibits MMPs that are involved in other molecular processes. Experiments, in which the alternative MMP inhibitor GM6001 (Illomastat, Galardin; **Wojtowicz-Praga et al., 1997**) was used, showed qualitatively similar results (not shown).

HB-EGF also functions as the cell surface receptor for diphtheria toxin and soluble HB-EGF can be sequestered by the non-toxic diphtheria toxin mutant CRM197 (**Mitamura et al., 1995**). Like Marimastat, CRM197 treatment (IP injections of 1 µg CRM197 6 h before and at the time of tissue damage) did not noticeably affect cell proliferation in the intact tissue but robustly reduced mitotic activity in response to tissue injury (**Figure 7B, E**). TrX treatment induced a 4.6 ± 0.7-fold increase in the number of BrdU positive cells in the core sensory OE of animals that were injected with the vehicle DMSO (**Figure 7F**; vehicle – ctr: 13.1 ± 2.0, TrX: 60.8 ± 9.6 cells/hemi OE; p_adj_ = 5.7×10^−9^, Tukey HSD *post-hoc* analysis on one-way ANOVA, F_(3,116)_ = 3.6×10^−11^). In contrast, only a nonsignificant 1.7 ± 0.2-fold increase in the number of cells could be detected in CRM197-treated fish (CRM197 – ctr: 9.8 ± 1.5, TrX: 22.1 ± 2.1 cells/hemi OE, p_adj_ = 0.320). Again, ongoing cell proliferation at the ILC was not affected by the presence of the inhibitor.

Ultimately, HB-EGF exerts its effect onto target cells through activation of EGFR, which can be blocked by the kinase inhibitor PD153035 (**Fry et al., 1994**). In these experiments, TrX lesions induced a 4.4 ± 0.3-fold increase in core sensory OE mitotic activity in animals injected with the vehicle DMSO (**Figure 7B, G, H**; vehicle – ctr: 8.8 ± 1.3, TrX: 38.8 ± 2.5 cells/hemi OE; p_adj_ = 5.0×10^−15^, Tukey HSD post-hoc analysis on one-way ANOVA, F_(3,116)_ = 78.9, p < 2.0×10^−16^). Even though tissue damage resulted in an increased number of proliferating cells in PD153035-treated animals (PD153035 – ctr: 4.1 ± 0.6, TrX: 16.5 ± 1.9 cells/hemi OE; p_adj_ = 9.5×10^−6^), the increase in mitotic activity was only 1.9 ± 0.2-fold higher in comparison to unlesioned vehicle controls (p_adj_ = 0.011). This is most likely due to the effect of the inhibitor on ongoing activity in the intact tissue, which was much lower in PD153035-injected animals. Taken together, these observations strongly suggest that HB-EGF/EGFR signaling is required during the early phase of the injury response to trigger HBC activity in the lesioned zebrafish OE at 24 hpl. Inhibition of HB-EGF/EGFR signaling prevents the development of the typical spatial and temporal pattern of mitotic activity that drives OSN neurogenesis and tissue regeneration. In contrast, inhibition has no major effect on GBC activity at the ILC and SNS, further strengthening the notion that HB-EGF/EGFR signaling is specifically involved in repair but not maintenance neurogenesis in the zebrafish OE.

### Inhibition of HB-EGF/EGFR signaling strongly diminishes OE tissue regeneration

The regenerative tissue response in the zebrafish OE comprises the early activation of HBCs at around 24 hpl, a peak in mitotic activity at around 72 hpl, and the recovery of about 80% of the OSN population by 120 hpl (**Kocagöz et al., 2022**). To further confirm the effect of HB-EGF/EGFR inhibition on OSN regeneration, we applied the pathway inhibitors around the time of the tissue lesion, labeled mitotic cells by incubation in BrdU between 48 and 72 hpl, and analyzed the BrdU and HuC/D staining pattern of intact and lesioned OEs at 120 hpl (**Figure 8**).

**Figure 8 –.**
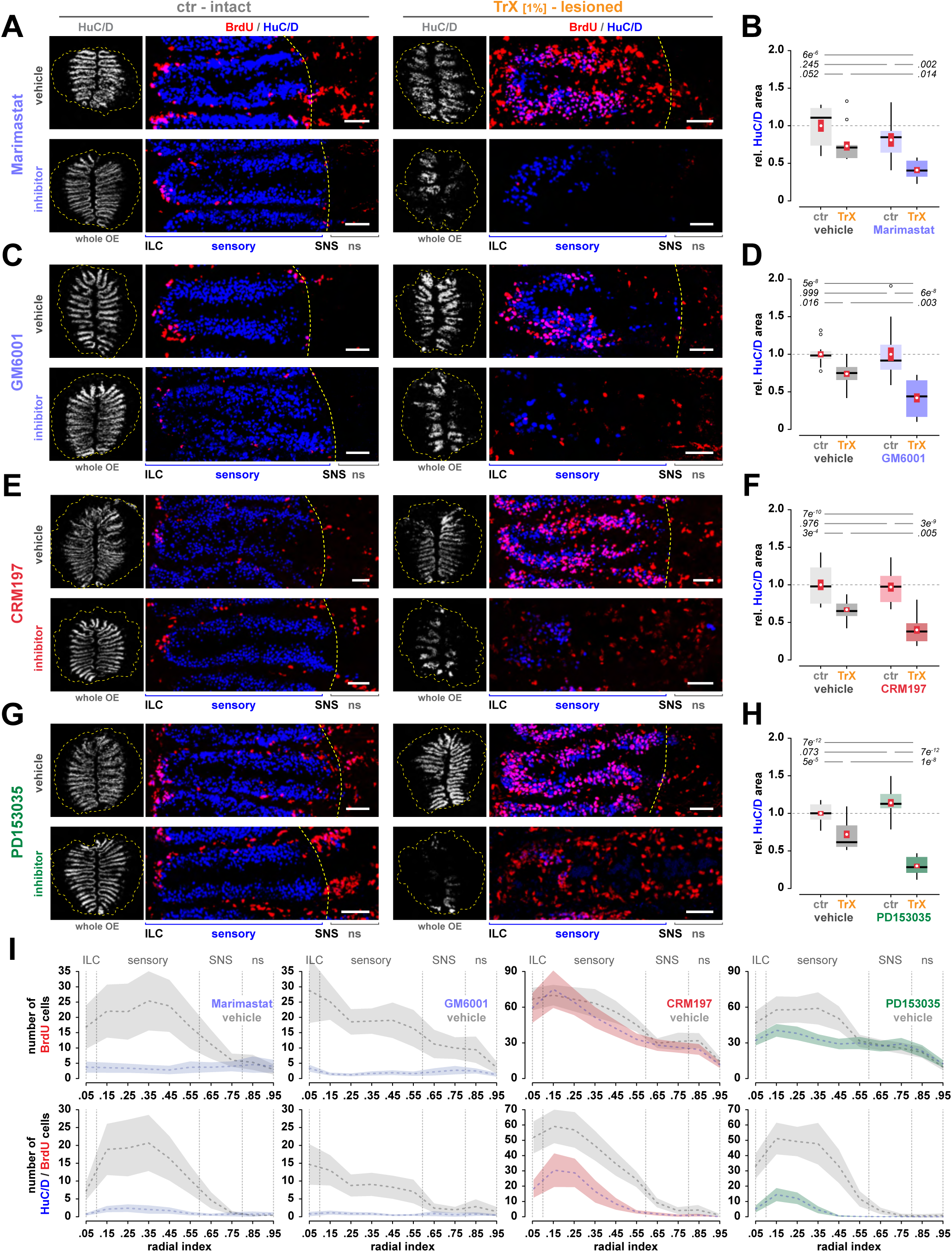
Inhibition of HB-EGF/EGFR signaling affects OSN regeneration. **A.** Immunohistochemistry against BrdU (red) and HuC/D (grey, blue) on intact control OEs (left) and OEs that have been lesioned with 1% TrX (right) of fish that were injected with vehicle (top panels) or the MMP inhibitor Marimastat (bottom panels). Fish were incubated in BrdU-containing water at the peak of injury-induced cell proliferation between 48 and 72 hpl and analyzed at 120 hpl. The small grey images on the left show the HuC/D staining pattern on full OE sections as analyzed for quantification of OSN regeneration; the yellow dashed line demarcates the outline of the tissue section. Color images show individual epithelial folds; interlamellar curves (ILCs) are to the left, the yellow dashed lines indicate the position of the sensory/non-sensory border (SNS). Note that injury-induced cell proliferation is severely reduced in Marimastat-treated fish and that the OE only partially regenerates from the lesion. Scale bars: 25 µm. **B.** Quantification of the efficiency of OE regeneration expressed as the tissue area covered by HuC/D-positive OSNs relative to the overall size of the OE section in vehicle- (grey, left) and Marimastat-injected fish (right, blue); ctr: unlesioned control OEs, TrX: OEs lesioned with 1% TrX. The HuC/D area has been normalized to the intact OEs of vehicle-treated animals. Control OEs regenerate 73.2 ± 6.3% of their OSNs by 120 hpl, while the HuC/D area is reduced to 41.0 ± 4.2% in Marimastat-treated specimens. The data comprise 10 tissue sections from 2 animals (5 sections/fish) for each condition; Box and Tukey-style whisker plots, the white dots and red vertical bars denote the means and SEM of the respective distributions, the p-values on the top indicate the result of a Tukey HSD *post-hoc* test on one-way ANOVA, F_(3,36)_ = 12.0, p = 1.2×10^−5^. Source data for the HuC/D area analysis are provided in **Figure 8 – Source data 1**. **C.** Immunohistochemistry against BrdU (red) and HuC/D (grey, blue) on intact and TrX-lesioned OEs of fish treated with the MMP inhibitor GM6001 (bottom panel) or vehicle (top panel). Same experimental setup as in A. Scale bars: 25 µm. **D.** Quantification of the HuC/D area in vehicle- (grey, left) and GM6001-treated fish (blue, right); same analysis as in B. Inhibition of MMPs by GM6001 reduces the efficiency of regeneration from 73.5 ± 4.0% to 41.9 ± 6.3%. The data comprise 15 tissue sections from 3 fish (5 sections/animal) for each condition. ANOVA: F_(3,56)_ = 20.5, p = 4.2×10^−9^. **E.** Immunohistochemistry against BrdU (red) and HuC/D (grey, blue) on intact and TrX-lesioned OEs of fish treated with the HB-EGF blocker CRM197 (bottom panel) or vehicle (top panel). Same experimental setup as in A. Scale bars: 25 µm. **F.** Quantification of the HuC/D area in vehicle- (grey, left) and CRM197-treated fish (red, right); same analysis as in B. Sequestration of HB-EGF by CRM197 reduces the efficiency of regeneration from 66.7 ± 3.2% to 40.3 ± 4.8%. The data comprise 15 tissue sections from 3 fish (5 sections/animal) for each condition. ANOVA: F_(3,56)_ = 27.6, p = 4.3×10^−11^. **G.** Immunohistochemistry against BrdU (red) and HuC/D (grey, blue) on intact and TrX-lesioned OEs of fish treated with the EGFR inhibitor PD153035 (bottom panel) or vehicle (top panel). Same experimental setup as in A. Scale bars: 25 µm. **H.** Quantification of the HuC/D area in vehicle- (grey, left) and PD153035-treated fish (green, right); same analyses as in B. Inhibition of EGFR kinase activity reduces the efficiency of regeneration from 71.6 ± 5.0% to 29.9 ± 3.0%. The data comprise 15 tissue sections from 3 fish (5 sections/animal) for each condition. ANOVA: F_(3,56)_ = 82.5, p <2×10^−16^. **I.** Radial tissue distribution of BrdU-positive cells (top row) and BrdU/HuC/D double-positive cells (bottom row) in lesioned OEs of vehicle-treated animals (grey) and fish injected with Marimastat (left, blue), GM6001 (middle left, blue), CRM197 (middle right, red), and PD153035 (right, green), which were incubated in BrdU-containing water between 48 and 72 hpl and analyzed at 120 hpl. The MMP inhibitors Marimastat and and GM6001 almost completely block injury-induced mitotic activity, while inhibition of soluble HB-EGF or EGFR activity results in delayed OSN generation as indicated by the reduction of double-positive cells despite increased cell proliferation at the same time point. The data represent the means and 95% confidence intervals of 20 (Marimastat) and 30 (GM6001, CRM197, PD153035) hemi-OEs from 2/3 fish (10 sections per animal) for each condition. Raw positional cell counts and statistics information (one-way ANOVA, Tukey HSD *post hoc* tests) for Marimastat-, GM6001-, CRM197-, and PD153035-treated fish are provided in **Figure 8 – source data 2 – 5**. **Figure 8 – Source data 1** **Source data for the effect of HB-EGF/EGFR inhibition on recovery of HuC/D-positive OSNS** This spreadsheet details the raw data and statistical analysis (one-way ANOVA, Tukey HSD *post hoc* tests) for the area covered by HuC/D-positive OSNs in Marimastat-, GM6001-, CRM197-, and PD153035-treated animals as shown in **Figure 8B, D, F**, and **H**. **Figure 8 – Source data 2** **Source data for the effect of inhibition of metalloprotease activity by Marimastat on injury-induced cell proliferation and OSN neurogenesis at 120 hpl** The spreadsheet provides the raw positional cell counts and statistics information (one-way ANOVA, Tukey HSD *post hoc* tests) for the number of BrdU and BrdU/HuC/D double-positive cells in Marimastat- and DMSO vehicle-treated fish. **Figure 8 – Source data 3** **Source data for the effect of inhibition of metalloprotease activity by GM6001 on injury-induced cell proliferation and OSN neurogenesis at 120 hpl** The spreadsheet provides the raw positional cell counts and statistics information (one-way ANOVA, Tukey HSD *post hoc* tests) for the number of BrdU and BrdU/HuC/D double-positive cells in GM6001- and DMSO vehicle-treated fish. **Figure 8 – Source data 4** **Source data for the effect of HB-EGF sequestration by CRM197 on injury-induced cell proliferation and OSN neurogenesis at 120 hpl** The spreadsheet provides the raw positional cell counts and statistics information (one-way ANOVA, Tukey HSD *post hoc* tests) for the number of BrdU and BrdU/HuC/D double-positive cells in CRM197- and DMSO vehicle-treated fish. **Figure 8 – Source data 5** **Source data for the effect of inhibition of EGFR activity by PD153035 on injury-induced cell proliferation and OSN neurogenesis at 120 hpl** The spreadsheet provides the raw positional cell counts and statistics information (one-way ANOVA, Tukey HSD *post hoc* tests) for the number of BrdU and BrdU/HuC/D double-positive cells in PD153035- and DMSO vehicle-treated fish.

Extended exposure to the MMP inhibitors Marimastat and GM6001 (IP injections at 4 h before the lesion, at the time of TrX treatment, 12 hpl, and 36 hpl) severely reduced mitotic activity and OSN generation (**Figure 8A, C**). Only few BrdU-positive cells could be detected in the core sensory region of lesioned OEs from inhibitor-treated animals at 120 hpl, while a strong upregulation of mitotic activity occurred in vehicle-treated control fish (Marimastat – DMSO: 71.0 ± 13.55, inhibitor: 9.25 ± 1.9 cells/hemi-OE, p_adj_ = 1.3×10^−7^, Tukey HSD *post-hoc* analysis on one-way ANOVA, F_(3,76)_ = 19.8, P = 1.4×10^−9^; GM6001 – DMSO: 56.4 ± 6.6, inhibitor: 4.8 ± 0.8 cells/hemi-OE, p_adj_ = 5.9×10^−14^, F_(3, 116)_ = 34.4, p = 5.5×10^−16^; **Figure 8I**, top panel). More importantly, the repopulation of the OE with HuC/D-positive OSNs was also severely reduced (**Figure 8B, D**). To quantify the effect on OE restoration, we thresholded the HuC/D signal, measured the area covered by HuC/D-positive OSNs relative to the size of the overall tissue section and normalized to unlesioned control OEs of vehicle-treated animals. While the lesioned OE of vehicle-treated control fish regained 73.2 ± 6.3% (Marimastat group: p_adj_ = 0.052, Tukey HSD *post-hoc* analysis on one-way ANOVA, F_(3,36)_ = 12.0, p = 1.3×10^−5^) and 73.5 ± 4.0% (GM6001 group: p_adj_ = 0.016, F_(3, 56)_ = 20.6, p = 4.2×10^−9^) of the HuC/D-positive area, this value was reduced to only 41.0 ± 4.2% (p_adj_ = 5.7×10^−6^) and 41.9 ± 6.3% (p_adj_ = 5.0×10^−8^) in Marimastat- and GM6001-injected fish, respectively. Thus, inhibition of MMPs results in less efficient OE regeneration and OSN restoration at 120h hpl.

A similar reduction in the restoration of HuC/D-positive OSNs could also be observed when HB-EGF and EGFR signaling were inhibited by CRM197 and PD153035, respectively (**Figure 8F, H**; CRM197: IP injection 4 h before TrX lesion, at the time of OE lesion, 24 hpl, and 48 hpl; PD153035: 4 h before lesion, concomitant with the lesion, and 4 hpl). While the HuC/D area recovered to 66.7 ± 3.2% (CRM197 group; p_adj_ = 2.8×10^−4^, F_(3, 56)_ = 27.6, p = 4.3×10^−11^) and 71.6 ± 5.0% (PD153035 group; p_adj_ = 5.1×10^−5^, F_(3, 56)_ = 82.5, p < 2×10^−16^) in vehicle-injected controls, only 40.3 ± 4.8% (p_adj_ = 7.1×10-10) and 29.9 ± 3.0% (_padj_ = 7.3×10^−12^) were reached in CRM197-and PD153035-treated fish, respectively. Surprisingly, however, the reduction in OE regeneration occurred despite increased proliferative activity in CRM197- and PD153035-treated fish following OE lesion (**Figure 8E, G, I**, top panel). In the lesioned OE of CRM197-injected animals, about 10-fold more mitotically active cells could be detected relative to intact control OEs (ctr: 15.0 ± 1.2, TrX: 157.9 ± 12.7 cells/hemi-OE; p_adj_ = 3.8×10^−14^, Tukey HSD *post-hoc* analysis on one-way ANOVA, F_(3,116)_ = 81.7, p < 2×10^−16^), which was similar to the 9-fold increase that occurred in vehicle-treated control animals (ctr: 21.2 ± 2.0, TrX: 186.0 ± 15.1 cells/hemi-OE, p_adj_ = 1.0×10^−14^). In PD153035-treted animals, the number of mitotic cells increased more than 3-fold (ctr: 29.2 ± 3.0, TrX: 99.8 ± 6.7 cells/hemi-OE; p_adj_ = 1.9×10^−6^, Tukey HSD *post-hoc* analysis on one-way ANOVA, F_(3,116)_ = 59.2, p < 2×10^−16^), which was less than in the control group (12-fold, ctr: 13.4 ± 1.7, TrX 167.4 ± 16.8 cells/hemi-OE, p_adj_ = 9.7×10^−15^) but much higher than for MMP inhibitors (**Figure 8I**).

We therefore suspected that OSN repair neurogenesis was severely delayed in CRM197- and PD153035-treated animals while it was completely blocked by Marimastat and GM6001. To follow up on this observation, we reanalyzed the changes in HuC/D/BrdU double-positive cells in TrX-lesioned OEs, which correspond to newly generated OSNs at 120 hpl (**Figure 8I**, bottom panel). While almost no double-positive cells could be detected in Marimastat- and GM6001-injected animals (Marimastat group – inhibitor: 9.1 ± 2.6, vehicle: 96.3 ± 18.2 cells/hemi-OE; p_adj_ = 1.5×10^−7^, F_(3, 76)_ = 16.6, P = 2.1×10^−8^; GM6001 group – inhibitor: 5.7 ± 1.5, vehicle: 68.1 ± 11.1 cells/hemi-OE, p_adj_ = 2.6×10^−10^, F_(3, 116)_ = 20.1, p = 1.4×10^−10^), OSNs were generated following CRM197 and PD153035 treatment. Nevertheless, a strong reduction in the number of newly generated OSNs was apparent for both inhibitor groups (CRM197 group (0.4-fold) - inhibitor: 111.1 ± 16.1, vehicle: 296.0 ± 26.8 cells/hemi-OE, p_adj_ = 5.9×10^−12^, F_(3, 116)_ = 49.7, p < 2×10^−16^; PD153035 group (0.2-fold) – inhibitor: 37.0 ± 5.4, vehicle: 232.0 ± 26.9 cells/hemi-OE, p_adj_ = 6.4×10^−14^, F_(3,116)_ = 40.6, p < 2×10^−16^). Thus, OSN neurogenesis was not completely blocked but severely delayed/reduced in the experimental regime that we applied. It remains to be explored whether the late induction of mitotic activity results from insufficient inhibition at the late analysis time point or the influence of redundant molecular pathways that stimulate delayed HBC activity.

## Discussion

Stem cell proliferation needs to be tightly controlled as either excessive or diminished activity would be detrimental to tissue function (**Clevers et al., 2014**). While exaggerated activity typically results in various forms of cancer, reduced activity would prevent the generation of a sufficient number of functional cells during growth, maintenance, and regeneration (**Clarke and Fuller, 2006**). HBCs constitute a neurogenic stem/progenitor cell population in the OE, which remains quiescent under physiological conditions but becomes selectively activated by tissue injury, while GBCs remain constitutively active during the life span of the organism to replenish sporadically dying OSNs (**Schwob et al., 2017**). Zebrafish HBCs strongly respond to the EGFR ligand HB-EGF (**Figure 5**), the expression of which is highly and transiently upregulated immediately after OE damage (**Figure 3**). *Hbegfa* reaches a peak of 14-fold higher expression by 4 hpl but declines to baseline expression shortly after 24 hpl (**Figure 1**). Exogenous stimulation of the OE with recombinant HB-EGF is sufficient to induce HBC expansion (**Figure 5**) and OSN neurogenesis (**Figure 4**), while inhibition of HB-EGF/EGFR signaling results in reduced HBC proliferation and delayed OSN restoration following injury (**Figure 6**, **7**). Thus, HB-EGF/EGFR signaling appears to be a critical component of the molecular signaling network that controls stem cell activity in zebrafish HBCs. In contrast, GBCs do not appear to be affected by HB-EGF and inhibition of HB-EGF/EGFR signaling in the intact tissue does not notably reduce their activity.

### Dual progenitor system and partially overlapping stem cell niches

GBCs and HBCs are both multipotent progenitor cells and essentially produce the same outcome, yet, do so under different tissue conditions. Both cell types have been shown to generate OSNs, in addition to SCs and gland/duct cells in the rodent OE (**Chen et al., 2004**; **Leung et al., 2007**), while their full lineage potential has not been addressed in zebrafish. The OE stem cell niche closely resembles other proliferating tissues with high regenerative capacity, such as skin and intestine, which are also endowed with a dual progenitor system (**Li and Clevers, 2010**). The occurrence of (and interconversion between) constitutively and conditionally active progenitor types has been suggested to be an adaptation of these tissues to prevent exhaustion of the available stem cell pool under different tissue conditions (**Visvader and Clevers, 2016**). Injury-activated HBCs generate a transient population of *ascl1*-expressing cells in mouse (**Leung et al., 2007**) and zebrafish (**Kocagöz et al. 2022**) that closely resemble GBCs. Conversely, the generation of HBCs from activated GBCs has also been suggested to occur in the mouse OE (**Lin et al., 2017**). Either way, it seems to be important that resident HBCs and GBCs respond selectively to tissue signals to fulfill their different functions. Selectivity could be achieved either by spatial segregation of the two cell types and differential exposure to niche factors or by inherent differences in their responsiveness to unique signals. Both cell types occupy the same spatial domain in the mammalian OE (**Schwob et al., 2017**), while zebrafish HBCs and GBCs overlap only at the ILC and SNS of the uninjured OE (**Demirler et al., 2020**). Under injury conditions, HBCs and HBC-derived GBC-like cells also overlap across the sensory region (**Kocagöz et al., 2022**), which exposes them to a similar molecular environment. Yet, differences in matrix attachment and cell-cell contacts, which further define their respective microenvironment (**Carter et al., 2004**) are likely to exist. Zebrafish GBCs have been shown to respond selectively to purine compounds that may be released from spontaneously dying OSNs, while HBCs do not (**Demirler et al., 2020**). In turn, we did not find indication that constitutive cell proliferation at the ILC and SNS are majorly affected by stimulation (**Figure 5**) or inhibition (**Figure 6**) of HB-EGF availability and EGFR signaling by CRM197 and PD153035, respectively. Thus, HB-EGF specifically, and EGFR signaling in general, appear not to provide direct inputs to resident GBCs at the ILC and SNS, while their potential influence on HBC-generated GBC-like cells has not yet been examined. In contrast, long-term exposure to MMP inhibitors strongly reduced proliferative activity at the ILC of the intact OE (not shown), suggesting that other MMP-sensitive signals provide input to resident GBCs.

### Possible interactions between HB-EGF/EGFR signaling and other signaling pathways

HB-EGF efficiently breaks HBC dormancy in the zebrafish OE (**Figure 4**). In rodents, however, a prominent role in HBC activation has been described for the Notch/Delta signaling pathway (**Herrick et al., 2017**). Cell-cell contacts in the intact OE keep Notch signaling high in non-dividing HBCs through interaction with the Notch ligand Jagged1 that is expressed by SCs. Notch activity, in turn, positively controls expression of the dormancy switch tp63 (**Packard et al., 2011**; **Fletcher et al., 2011)**. Both Notch and tp63 become downregulated when contacts between HBCs and SCs are disrupted as a consequence of tissue injury (**Herrick et al., 2017**). Loss of tp63 expression is necessary and sufficient in the mouse OE to induce HBC proliferation (**Schnittke et al., 2015**), which ultimately requires activation of Wnt signaling in rodents (**Wang et al., 2011**; **Fletcher et al., 2017**) and zebrafish (**Kocagöz et al., 2022**). Given that stimulation of EGFR activity by HB-EGF (this study) or other ligands (**Mahanthappa and Schwarting, 1993**; **Farbman and Buchholz, 1996**; **Getchell et al., 2000**; **Chen et al., 2020**) also robustly induces mitotic activity in HBCs suggests that regulation of HBC activity is more complex and that redundant pathways or synergistic interactions between diverse signaling routes may converge to drive Wnt activation and HBC expansion.

Mutual interactions between EGFR, Notch, and Wnt signaling are well documented for a variety of cancers and developing or regenerating tissues (**Sundaram, 2005**; **Doroquez and Rebay, 2006**; **Baker et al., 2014**; **Acar et al., 2021**). More specifically, crosstalk between HB-EGF-mediated EGFR activity and Notch signaling in the nervous system has been demonstrated for the neurogenic niche of the subventricular zone (**Aguirre et al., 2010**), for retinal progenitor cells (**Wan et al., 2012**; **Angbohang et al., 2014**), and for proliferating astrocytes *in vitro* (**Pushmann et al., 2014**). EGFR signaling can reduce Notch pathway activity either by directly inhibiting receptor expression at the transcriptional level (**Kolev et al., 2008**) or by antagonizing downstream pathway mediators. In the subventricular zone, EGFR activity promotes neural stem cell proliferation by upregulating the modifier protein NUMB to reduce levels of the Notch intracellular signaling domain (**Aguirre et al., 2010**), while phosphorylation of the downstream mediator and transcriptional co-repressor Groucho (Transducin-like enhancer protein; groucho-related gene in mammals) has been implicated in a number of developmental processes (**Hasson and Paroush, 2006**). Thus, EGFR stimulation by HB-EGF could potentially modulate Wnt signaling in HBCs either directly, by activating the Wnt effector β-catenin through a non-canonical route (**Knight et al., 2019**), or indirectly, by reducing Notch activity (**Acar et al., 2021**), similar to HBC/SC contact disruption in the rodent OE.

Inhibition of MMPs had stronger and longer lasting effects on injury-induced cell proliferation and OSN regeneration than direct sequestration of HB-EGF by CRM197 or inhibition of EGFR phosphorylation by PD153035 (**Figure 7**). This is most likely because the MMP inhibitors Marimastat and GM6001 have wide target ranges, which include MMPs that are relevant for other signaling pathways (**Wojtowicz-Praga et al., 1997**; **Löffek et al., 2011**). Interestingly, MMPs and ADAMs have also been shown to contribute to cleavage of the extracellular domains of Notch receptors and ligands (**Groot and Vooijs, 2012**). Thus, MMP inhibitors may affect Notch signaling in addition to HB-EGF shedding to amplify their effect on HBCs. In turn, Notch signaling can influence MMP expression and HB-EGF shedding (**Zhou et al., 2012**; **Diaz et al., 2013**), confounding the range of possible interactions between these pathways. CRM197 and PD153035 strongly prevented early (24 hpl; **Figure 6**) but not late (120 hpl; **Figure 7**) induction of injury-induced cell proliferation. A potential reason may be that HB-EGF/EGFR inhibition did not persist over the extended duration of the experiment due to limitations in the half-life of the inhibitors. Interestingly, similar observations were made for Illomastat (GM6001) and the EGFR inhibitor gefinitib in the mouse OE, where both inhibitors severely reduced but did not fully prevent injury-induced HBC expansion (**Chen et al., 2020**). Therefore, we cannot exclude the possibility that signaling through redundant pathways, such as Notch signaling, and their convergence on Wnt activity resulted in delayed HBC activation. In contrast, inhibition of MMPs completely abolished cell proliferation and OSN neurogenesis even at 120 hpl, which suggests that MMP activity is essential during an early and narrow time window and that HBC activation only ensues when MMP activity is available shortly after tissue damage.

The downregulation of tp63 by loss of Notch signaling during activation of rodent HBCs is unusual because activated, not reduced Notch signaling tunes down tp63 expression in other proliferating tissues (**Nguyen et al., 2006**; **Tadeu and Horsley, 2013**). Surprisingly, tp63 is highly upregulated in the injured zebrafish (**Supplementary Figure 1**) and mouse OE (**Chen et al., 2020**). Levels of tp63 expression rise 4.0 ± 0.1-fold by 4 hpl in zebrafish and remain significantly elevated until 72 hpl. Our attempts to follow tp63 expression in Krt5-positive HBCs at tissue level also did not reveal indication for downregulation (not shown). Rather, an increased number of tp63-positive cells emerge after the lesion, suggesting that tp63 does not function as a dormancy switch in zebrafish HBCs. Alternative cellular signaling routes, such as activation of MAP kinase and PI3K/AKT signaling downstream of EGFR, synergistically or redundantly to Notch, may play a more prominent role through direct induction of Wnt signaling (**Hu and Li, 2010**; **Knight et al., 2019**; **Acar et al., 2021**) in zebrafish HBCs.

### HB-EGF and other EGFR ligands

Other EGFR ligands besides HB-EGF have been documented to promote HBC proliferation *in vivo* (**Farbman and Ezeh, 2000**; **Getchell et al., 2000**) or *in vitro* (**Mahanthappa and Schwarting, 1993**; **Farbman and Buchholz, 1996**). EGF, TGF-α, and amphiregulin stimulate mitogenic activity in the native OE or cultured OE cells. For instance, overexpression of TGF-α in HBCs under control of the *Krt14* promoter that is active in HBCs stimulates 6-fold higher HBC proliferation but does not affect GBC activity (**Getchell et al., 2000**). We show that zebrafish HBCs, like their rodent counterparts, express EGFR (**Figure 2**; **Holbrook et al., 1995**; **Krishna et al., 1996**; **Chen et al., 2020**), and are most likely direct targets of activated EGFR ligands. However, despite their prominent mitosis-promoting effect on HBCs, upregulated expression of canonical EGFR ligands has previously not been validated in the injured OE of any species. TGF-α is expressed prominently by Bowman’s gland cells in rodents and less strongly by basal and supporting cells (**Farbman and Buchholz, 1996**). Yet, neither TGF-α, EGF, nor amphiregulin appear to be upregulated in a single cell transcriptomics study of the methimazole-injured mouse OE (**Gadye et al., 2017**). Recently the L1 cell adhesion molecule family member NrCAM has been implicated in HBC activation through an EGFR-dependent mechanism (**Chen et al., 2020**). Even though NrCAM is not a classical EGFR ligand, it has been shown to modulate EGFR activity (**Zhang et al., 2017**; **Chen et al., 2020**). Interestingly, the zebrafish homolog *nrcama* is downregulated by 40 – 60% between 4 and 24 hpl in the lesioned OE (**Figure 1B**), while *Hbegf* is also upregulated in the injured mouse tissue (**Gadye et al., 2017**), suggesting that HB-EGF may be a biologically more relevant EGFR ligand in both species. In as much HB-EGF and NrCAM synergize in the mouse OE awaits further investigation.

### HB-EGF expression, self-amplification and gene regulatory networks

HB-EGF is expressed in a low number of cells in the intact zebrafish OE, most of which stain positive for the basal cell marker Sox2 or the HBC marker Krt5 (**Figure 3**). Expression of *hbegfa* is upregulated in response to tissue damage in all major OE cell types, comprising OSNs, SCs, and basal progenitors. The mechanisms regulating this broad induction of HB-EGF expression in the injured OE is elusive. In the zebrafish retina, HB-EGF is part of a complex gene regulatory network that comprises other growth factors, insulin, and cytokines (**Wan et al., 2014**). Stimulation with any of these factors prompted mutual upregulation, thereby, promoting a bistable network switch between OFF and ON states (**Nagata and Kikuchi, 2020**). Activation of EGFR by HB-EGF triggers additional positive feedback loops, which include the auto-/paracrine stimulation of further HB-EGF shedding and upregulation of *Hbegf* expression, to ensure robust EGFR activation in target cells (**Hashimoto et al., 1994**; **Umata et al., 2001**). The observations that exposure to exogenously applied recombinant HB-EGF induces strong *hbegfa* transcription in HBCs (**Figure 5H, I**) and that tissue damage upregulates EGFR expression (**Figure 1A**) further support this view. Thus, tissue- or immune cell-derived cytokines may trigger *hbegfa* expression in the injured zebrafish OE, which further amplifies *hbegfa* expression by HBCs in a process that is equivalent to findings in the retina (**Wan et al., 2012**; **Zhao et al., 2014**). The functional implication of such a mechanism could be that the system becomes robust against noise to prevent precocious activation but retains the potential to respond once a meaningful threshold is exceeded. We employed an experimental injury model that globally destroys OE cells and, accordingly, observed widespread induction of *hbegfa* expression (**Figure 3**) and activation of HBC proliferation (**Figure 6**). However, localized microlesions would be expected to occur more likely under natural conditions. Paracrine stimulation between neighboring cells may result in recruitment of additional HBCs and, thus, a wider footprint of activated stem cells to repair the injury more efficiently. Support for such a scenario comes from focal lesion experiments in the zebrafish retina, where inhibition of ectodomain shedding and EGFR signaling restricted the domain of HB-EGF expression and progenitor cell proliferation to the injured site, while more distant cells also responded to the injury when no inhibitor was applied (**Wan et al., 2012**).

### Does HB-EGF promote proliferation, differentiation, or both?

EGFR signaling has been shown to promote cell proliferation (**Wee and Wang, 2017**) but also cell/lineage differentiation (**Castanieto et al., 2015**) and cell survival by inhibition of apoptosis (**She et al., 2005**). The relatively wide range of effects is linked to the manifold intracellular signaling pathways that can be stimulated downstream of ErbB receptors, which include the MAP kinase, PI3K-AKT, SRC, PLC-γ1-PKC, JNK, and JAK-STAT pathways, in addition to endocytosis and direct nuclear translocation of receptor complexes (**Wee and Wang, 2017**). Of those, MAP kinase and AKT signaling activate mitotic cell divisions, largely through regulation of cyclin D (**Pennock and Wang, 2003**), while AKT also inhibits apoptosis by inhibiting BAD (**She et al., 2005**) and caspase 9 (**Cardone et al., 1998**).

Proliferation and differentiation often show an antagonistic relationship and cell cycle-promoting signals must wane before differentiation can ensue (**Ruijtenberg and van den Heuvel, 2016**). Exogenous stimulation of the OE with HB-EGF promotes strong HBC proliferation (**Figure 4**), suggesting that EGFR signaling controls exit from cell cycle arrest in quiescent HBCs. However, this observation does not inform whether HB-EGF-activated HBCs only undergo amplification of the stem cell pool through symmetric/self-renewing divisions or also differentiation by dividing asymmetrically (**Casas Gimeno and Paridaen, 2022**). HB-EGF stimulation increases the number of *ascl1*-positive GBCs (**Figure 5G**) and OSN neurogenesis (**Figure 4D**), thus, at least some of the newly generated cells have the potential to differentiate and to enter the neurogenic lineage. It has been shown that EGFR signaling can exert symmetry-breaking effects in some contexts (**Ayuso-Sacido et al., 2010**; **Wang et al., 2019**) but it remains elusive whether activated HBCs depend on HB-EGF/EGFR signaling to commit to a neuronal fate. Thus, it is possible that HBCs display inherent population asymmetry (**Simons and Clevers, 2011**) and that some of their daughter cells either differentiate spontaneously or under the influence of additional factors that are either ubiquitously present in the HBC niche or induced as a consequence of the injury. We have previously shown that HBCs can spontaneously give rise to neurogenic clones in the sensory OE (**Kocagöz et al., 2022**), albeit at low frequency. Enhanced HBC proliferation in HB-EGF-stimulated OEs may increase the number of HBCs that undergo spontaneous differentiation, which may account for the observed neurogenic activity in response to HB-EGF stimulation even in the absence of an injury.

## Conclusion

Expression of EGFR by injury-responsive HBCs has been demonstrated more than 25 years ago (**Holbrook et al., 1995**; **Krishna et al., 1996**), however, the identification of biologically relevant ligands has never been reported. To our knowledge, HB-EGF is the first functionally identified canonical EGFR ligand that is naturally expressed and upregulated in the injured OE. Induction of HB-EGF expression precedes HBC expansion and may, therefore, constitute an important early molecular signal to break HBC dormancy, which sets in motion a cascade of cell proliferation and differentiation events that results in repopulation of the injured OE with OSNs. HB-EGF is sufficient to induce HBC proliferation and OSN neurogenesis and necessary for full and timely recovery of the tissue from injury. OEs, in which either HB-EGF or EGFR function has been inhibited show a significant delay and reduction in the efficiency with which OSNs are generated following the lesion. HB-EGF/EGFR signaling may interact with other signaling pathways that have synergistic functions or independently converge onto mitosis-promoting cell cycle regulators, suggesting the presence of a complex and interconnected regulatory signaling network that controls HBC activity.

## Material and Methods

### Zebrafish maintenance and husbandry

Adult zebrafish (*Danio rerio*; ≥ 6 months of age) of the AB/AB laboratory strain (RRID:ZIRC_ZL62) and an in-house line derived from a local pet shop were used for experiments. Fish were bred, grown, and kept at the AAALAC-accredited research animal facility (Vivarium) of Boğaziçi University, Center for Life Sciences and Technologies (Istanbul) and maintained in commercial housing systems (Aquatic Habitats, FL, USA) at constant temperature of 27°C and under a 14 h light/10 h dark regime. Artificial fresh water (2.0 g sea salt, 7.5 g sodium bicarbonate, and 0.84 g calcium sulfate dissolved in 100 l of reverse osmosis water) was resupplied daily. Fish were fed twice daily with flake food (TetraMin; Tetra, Melle, Germany) supplemented once per day with live brine shrimp larvae (*Artemia sp.*; Fides Artemia, Istanbul, Turkey) or frozen food cubes containing a mixture of brine shrimp, insect larvae, and algae. Fish were anesthetized during experimental interventions, such as lesioning of the OE or intraperitoneal injection (IP) of pharmacological agents, by submersion in 160 mg/l MS-222 (ethyl 3-aminobenzoate methanesulfonate; Sigma-Aldrich, Merck, Darmstadt, Germany) dissolved in tank water. The use of zebrafish for this study was approved by the Institutional Ethics Board for Animal Experiments at Bogazici University (BÜHADYEK) under title “The role of heparin-binding epidermal growth factor (HB-EGF) signaling during regenerative neurogenesis in the zebrafish olfactory epithelium”; approval date: 07.09.2018, amendment date: 26.08.2020. All experiments and procedures were performed in accordance with relevant guidelines and regulations pursuant to the use of animals in biological research, including the National Animal Protection Act (Turkish national law number 5199, “Hayvanları Koruma Kanunu”, published 24.06.2004 and amendment published 14.07.2021), the directive 2010/63/EU of the “European Parliament and the Council of 22. September 2010 on the Protection of Animals Used for Scientific Purposes” and the “Guide for the Care and Use of Laboratory Animals” (NRC2011) of the Association for Assessment and Accreditation of Laboratory Animal Care (AAALAC). All experimental manipulations on live animals were performed under MS-222 anesthesia and every effort was made to minimize suffering.

### Generation of OE lesions by irrigation with Triton X-100

To generate experimental lesions of the OE, adult fish (≥ 6 months of age) were anesthetized by submersion in 160 mg/l MS-222 and mounted between wet sponges onto the stage of a stereomicroscope (Zeiss; Jena, Germany). Approximately 1 - 2 µl of solutions containing 0.1 or 1% of the non-ionic detergent Triton X-100 (TrX; Sigma-Aldrich), dissolved in 0.1 M phosphate-buffered saline (PBS; 137 mM NaCl, 2.7 mM KCl, 10 mM Na_2_HPO4, 1.8 mM KH_2_PO_4_), supplemented with 0.1% phenol red (Sigma-Aldrich), were injected into one of the paired olfactory cavities through fine glass capillaries using a pressurized (Eppendorf FemtoJet Express; Eppendorf, Hamburg, Germany) or hydraulic (IM-6; Narishige, Tokyo, Japan) injection system. Care was taken to avoid spillover of the TrX solution to the internal control OE on the other side of the head. TrX solutions were injected twice at 45 s intervals for a total exposure time of 90 s. The detergent solution was flushed out from the nasal cavity at the end of the incubation period by a gentle stream of freshwater delivered from a Pasteur pipette before the animals were placed back into recovery tanks. The effect of TrX damage was compared either between lesioned and unlesioned control OEs of the same animal or across age-matched animals from the same breeding cycle (within one month of weekly breeding) when groups of animals were analyzed for the effect of pharmacological intervention.

### Labeling of mitotic activity by incorporation of thymidine analogues

To visualize proliferating cells, fish were incubated in freshly prepared tank water containing 30 mg/l of the halogenated thymidine analog 5-bromo-2’-deoxyuridine (BrdU; Sigma-Aldrich) for periods between 12 h and 24 h at various time points following experimental intervention and kept in the dark at 28°C. Label-positive mitotically active cells were detected by immunohistochemistry against BrdU on OE tissue sections. In experiments in which 5-ethynyl-2’-deoxyuridine (EdU; Sigma-Aldrich) was used, fish were injected with 30 µl of 10 mM EdU suspended in PBS and visualized using the Click-iT™ EdU Cell Proliferation Kit (Thermo Fisher Scientific, Waltham, MA, USA) according to the manufacturer’s instructions.

### Immunohistochemistry

For standard immunohistochemical detection of cell type-specific markers and mitotic labels, OEs were dissected in ice-cold 1x PBS and mounted in Tissue-Plus O.C.T. Compound (Thermo Fisher Scientific). Sections of 12 μm thickness were taken on a CM3050S cryostat (Leica; Wetzlar, Germany), dried at 65°C in a hybridization oven for ≥ 60 min, fixed with 4% paraformaldehyde (PFA; Sigma-Aldrich; dissolved in 1x PBS, pH = 7.4) for 15 min at room temperature (RT), and subsequently washed three times with PBS containing 0.1% (v/v) Tween 20 (Sigma-Aldrich; PBST) for 10 min each. For detection of the thymidine analog BrdU, sections were additionally incubated in 4N HCl for 15 min at RT for permeabilization of nuclei and washed 3 times with PBST for 10 min each. Blocking of unspecific antigens was carried out by incubation in 3% bovine serum albumin (BSA; Sigma-Aldrich; dissolved in PBST) for 1 h at RT. Tissue sections were incubated overnight at 4°C in a humidified chamber with primary antibodies: rat anti-BrdU (1:500; Cat#: ab6236, RRID:AB_305426; Abcam, Cambridge, UK), mouse anti-HuC/D (1:500; Cat#: A-21271, RRID:AB_221448; Thermo Fisher Scientific), rabbit anti-Krt5 (1:500; Cat#: 905501, RRID:AB_2565050; BioLegend, San Diego, CA, USA), rabbit anti-p63 (1:500; Cat#: GTX124660, RRID:AB_11175363; Genetex, Irvine, CA, USA), mouse anti-cytokeratin type II (1:1.000; Cat#: 1h5, RRID:AB_528323; Developmental Studies Hybridoma Bank, Iowa City, IA,USA), followed by three washes in PBST for 15 min each. For fluorescent detection, Alexa 488-, Alexa 555-, and Alexa 647-coupled anti-mouse, anti-rabbit, and anti-rat antibodies (1:800; Cat#: A-11001, RRID:AB_2534069; Cat#: A-21422, RRID:AB_141822; Cat#: A-21236, RRID:AB_2535805; Cat#: A-21434, RRID:AB_141733; Cat#: A-21247, RRID:AB_141778; Cat#: A-11034, RRID:AB_2576217; Cat#: A-21428, RRID:AB_2535849; Thermo Fisher Scientific), or Cy2-conjugated anti-mouse and Cy5-conjugated anti-rat IgGs (1:200; Cat#: 115-225-146, RRID:AB_2307343; Cat#: 112-175-143, RRID:AB_2338263; Jackson ImmunoResearch, Cambridge, UK) were used. Sections were incubated for 2 h at RT in secondary antibody solution, subjected to three successive washes in PBS for 10 min each, and visualized by confocal microscopy.

For detection of basal cell populations using a rabbit anti-Sox2 antibody (1:500; polyclonal, Cat#: GTX124477, RRID:AB_11178063; Genetex), a sodium citrate buffer-based antigen retrieval protocol was applied. OEs were treated with 4% PFA overnight at 4°C immediately after dissection, subsequently washed three times with PBST for 15 min each at 4°C, and mounted in Tissue-Plus O.C.T. Compound for cryosectioning. Tissue sections were air-dried overnight at RT and incubated in pre-heated sodium citrate buffer (10 mM Na_3_C_6_H_5_O_7_, 0.05% Tween 20, pH: 6.0) just below 100°C for 40 min. Sections were then placed in 1x PBS to cool down to RT for 10 min before applying the standard staining protocol starting from the HCl treatment step. In order to increase the quality of staining, primary antibody incubation was carried out for 48 h at 4°C and secondary antibody incubation was extended to 24 hours at 4°C.

### *In situ*-hybridization

*In situ*-hybridization against *hbegfa*, *egfr*, and *ascl1a* mRNA transcripts were performed as described previously (**Bayramli et al., 2017**). OE tissue sections were fixed in 4% PFA for 10 min, washed in 1x PBS treated with diethyl pyrocarbonate (PBS_DT_; Sigma-Aldrich), and incubated with 0.05% (v/v) proteinase K (Roche, Merck, Darmstadt, Germany) dissolved in 0.1 M Tris-HCl (pH 8.0) at 37 °C for 7.5 min. Sections were re-fixed in 4% PFA for 5 min and incubated in 0.2 M HCl for 10 min at RT before treatment with tri-ethanolamine buffer (662.5 µl tri-ethanolamine, 112 µl 1 M HCl, 49.1 µl H_2_O_DT_, 125 µl acetic anhydride; Roche). Slides were preincubated at 65°C in hybridization buffer (50% formamide (Sigma-Aldrich), 5x sodium-sodium citrate (Sigma-Aldrich), 50 µg/ml heparin (Sigma-Aldrich), 500 µg/ml yeast RNA (Sigma-Aldrich), 9.2 mM citric acid, 0.05% Tween 20) before exchange of the pre-incubation solution for hybridization buffer containing 3 ng/µl of digoxigenin-labeled (Roche) riboprobe. The riboprobes correspond to antisense transcripts of the zebrafish *hbegfa* (821 nt fragment of ENSDART00000109138.4; positions 347 – 1.168), *egfra* (449 nt fragment of ENSDART00000164152.3; positions 3.615 – 4.064), and the *ascl1a* (422 nt fragment of ENSDART00000056005.5; positions 638 – 1060; **Kocagöz et al., 2022**) genes that were amplified from zebrafish cDNA using specific primer pairs. The hybridized slides were washed in descending concentrations of sodium-sodium citrate (SSC) buffer (5x SSC, 2x SSC/50% formamide, 2x SSC, 0.2x SSC) at the hybridization temperature. The bound ribopropbe was detected using an alkaline phosphatase-conjugated goat anti-DIG Fab fragment (1:750; Cat#: 11093274910, RRID:AB_514497; Roche) and the HNPP/FastRed detection kit (Roche). Immunohistochemistry against cell type-specific markers and detection of proliferating cells was carried out subsequently to HNPP detection.

### Stimulation of the OE with recombinant HB-EGF

For direct stimulation of the OE with recombinant HB-EGF, fish were briefly anesthetized with 160 mg/mL MS222 in tank water, placed between wet sponge material onto the stage of a stereomicroscope, and kept anesthetized for 30 min with a gentle stream of 80 mg/L MS222 delivered to the mouth from a continuous perfusion system. Approximately 1 μl of solution containing human recombinant HB-EGF (RD Systems, Minneapolis, MN, USA) dissolved in 0.1% BSA/1x PBS was injected into one nasal cavity using a hydraulic injection system (IM-6; Narishige, Tokyo, Japan), while the contralateral cavity was injected with an equal volume of vehicle to serve as internal control. HB-EGF and vehicle solutions were resupplied three times at 10 min intervals to avoid drying. For the establishment of dose-response relationships, OEs were stimulated twice at 24 h intervals with PBS, 20, 100, or 200 ng/µl HB-EGF before labeling of mitotic activity by BrdU or EdU incorporation for 24 h. Animals that were examined for the induction of mitotic cells and immunohistochemistry against Krt5 and Sox2 or for the induction of *ascl1a* expression were stimulated twice with 200 ng/µl recombinant protein before dissection at 24 h following the last injection. The effect of HB-EGF stimulation on OSN neurogenesis was examined by incubation in BrdU for the first 12 h and analysis at 72 h post stimulation. Fish that were analyzed for *hbegfa* expression received a single injection of 200 ng/µl HB-EGF and were dissected 4 h post stimulation.

### Pharmacological inhibition of HB-EGF/EGFR signaling

To prevent HB-EGF ectodomain shedding, the broad-spectrum metalloprotease inhibitors Marimastat (CAS: 154039-60-8; Sigma-Aldrich) and GM6001 (Illomastat; CAS: 142880-36-2; Tocris, Bristol, UK) were used. For the analysis of short-term effects of metalloprotease inhibition on injury-induced cell proliferation at 24 hpl, fish received two priming IP injections of 80 µg/g Marimastat (dissolved in 10% DMSO/1x PBS; 50 μl injection volume) at 12 and 24 h before induction of OE lesions by nasal irrigation with 1% TrX, concomitant with the TrX lesion, and at 12 hpl. For the analysis of the effect of metalloprotease inhibition on OE regeneration by HuC/D immunohistochemistry, fish received a priming injection of 80 µg/g Marimastat or 50 µg/g GM6001 (dissolved in 25% DMSO/1x PBS) 4 h before the lesion, in addition to injections at the time of TrX treatment, 12 hpl, and 36 hpl. Soluble HB-EGF was sequestered using the diphtheria toxin mutant CRM197 (CAS: 92092-36-9; Santa Cruz Biotechnology, Dallas, TX, USA) resuspended in 100 mM HEPES Buffer. For the examination of early effects, animals received IP injections of 1 μg/g CRM197 in 30 μl PBS at 6 h before nasal irrigation with TrX and at the time of lesion. For analysis of the late effect of CRM197 on OSN regeneration, fish received IP injections at 4 h before, at the time of OE lesion, 24 hpl, and 48 hpl. Inhibition of EGFR signaling was performed by injecting 80 μg/g of the kinase inhibitor PD153035 (CAS: 183322-45-4; Sigma-Aldrich), dissolved in DMSO/1xPBS, at 4 h before TrX lesion, at the time of lesion and 4 h after the lesion for both short-term and long-term analysis. Independent control animals were injected with an equal volume of vehicle and the same time regime for each inhibitor. Fish were incubated in BrdU-containing tank water immediately after the lesion until analysis of early effects at 24 hpl, or between 48 and 72 hpl for analysis of late effects at 120 hpl.

### Transcriptome profiling and analysis

For transcriptome profiling, a previously generated data set (**Kocagöz et al., 2022**) comprising RNA-sequencing data for untreated control OEs and OEs dissected at 4, 12, 24, 72, and 120 hpl was augmented with an independent third biological replicate. Bioinformatics analysis was performed as described previously (**Kocagöz et al., 2022**) and read counts were converted to fragments per kilobase of transcript per million reads mapped (FPKM) using the countToFPKM package (**Alhendi, 2019**) in R (**R Core Team 2022**) and gene/genome information obtained from the Ensembl data base (https://www.ensembl.org/Danio_rerio/Info/Index; **Cunningham et al., 2022**) using the BioMart tool (**Smedley et al., 2009**). FPKM data were further processed in R for extracted genes of interest.

### Imaging and image processing

Stained tissue sections were visualized as confocal z-stacks on Leica SP5-AOBS or Leica SP8 confocal microscopes (Leica Microsystems, Wetzlar, Germany) at 1.024×1.024 and 2.048×2.048 pixels resolution using 20x and 40x water immersion lenses. Image stacks were further processed using Leica-LAS-AF and Fiji software (**Schindelin et al. 2012**) to generate maximum projections, to apply suitable look-up tables, to adjust brightness and contrast, and to crop relevant image regions for analysis and visualization purposes. No nonlinear image adjustments (e.g. gamma), or selective removal, combinations, and repositioning of image features were applied.

### Quantitative measurements and statistical evaluation

Counts and positional profiling of marker-positive cells (e.g. cells labeled with thymidine analogs or by immunohistochemistry) were performed as described previously (**Demirler et al., 2020**; **Kocagöz et al., 2022**). Briefly, individual epithelial folds or hemi-OEs were cropped from projected z-stacks, oriented horizontally, and divided into ten equidistant bins between the ILC and the peripheral margin of the tissue section. Hemi-OEs were selected between the ILC and the peripheral border with a 3×2 horizontal to vertical aspect ratio centered onto the horizontal midline of the tissue section. Left hemi-OEs were flipped horizontally before counting. Marker-positive cells were identified by supervised thresholding or manual tagging in Fiji. Cells were counted into bins using customized Fiji macros and further processed to corrected for cells that fall onto the borders of compartments. For the analysis of OE regeneration following lesions, the area covered by HuC/D positive cells was analyzed by measuring signals above threshold and dividing by the overall area of the tissue section as described previously (**Kocagöz et al., 2022**). Values were normalized to intact control OEs of vehicle-treated animals. Quantitative data were statistically analyzed and graphed in R (**R Core Team 2022**) using standard functions (boxplot, points, lines, polygons, etc.) and further formatted. One-way analysis of variance (ANOVA) and *post hoc* Tukey honestly significant difference (Tukey HSD) and Dennett’s tests were performed in R or IBM SPSS Statistics version 25.

## Acknowledgments

Work on this study was supported by The Scientific and Technological Research Council of Turkey (TÜBITAK) Grant Number 119Z081 to SHF. YK acknowledges support from TÜBITAK-BIDEB 2211-A. The authors are grateful to Umut Sahin for valuable suggestions and reagents.

## Author contributions

Investigation: SS, YK, ASA, KG, ZD, EB, MCD, SHF; Visualization: SS, YK, ASA, KG, ZD, SHF; Formal Analysis: SS, YK, ASA, ZD, EB, MCD, SHF; Writing – Original Draft Preparation: SHF; Writing – Review & Editing: SS, YK, ASA, KG, ZD, EB, MCD, SHF: Conceptualization, Supervision, Project Administration, and Funding Acquisition: SHF.

## Data availability

All data generated or analyzed during this study are included in the manuscript and supporting files; Source Data files are provided for Figures 1, 2, 4, 5, 6, 7, and 8.

## References

1. Abud HE, Chan WH, Jardé T (2021) Source and Impact of the EGF Family of Ligands on Intestinal Stem Cells Frontiers in Cell and Developmental Biology 9:685665. https://doi.org/10.3389/fcell.2021.685665

2. Acar A, Hidalgo-Sastre A, Leverentz MK, Mills CG, Woodcock S, Baron M, Collu GM, Brennan K (2021) Inhibition of Wnt signalling by Notch via two distinct mechanisms Scientifc Reports 11:9096. https://doi.org/10.1038/s41598-021-88618-5

3. Aguirre A, Rubio ME, Gallo V (2010) Notch and EGFR pathway interaction regulates neural stem cell number and self-renewal Nature 467:323–7. https://doi.org/10.1038/nature09347

4. Alhendi A (2019) countToFPKM: Convert Counts to Fragments per Kilobase of Transcript per Million (FPKM) https://CRAN.R-project.org/package=countToFPKM

5. Angbohang A, Eastlake K, Becker S, Wu N, Limb GA (2014) Upregulation of the Notch and Wnt signalling pathways by HB-EGF in adult human Müller stem cells in vitro Investigative Ophthalmology & Visual Science 55:1371

6. Asakura M, Kitakaze M, Takashima S, Liao Y, Ishikura F, Yoshinaka T, Ohmoto H, Node K, Yoshino K, Ishiguro H, Asanuma H, Sanada S, Matsumura Y, Takeda H, Beppu S, Tada M, Hori M, Higashiyama S (2002) Cardiac hypertrophy is inhibited by antagonism of ADAM12 processing of HB-EGF: metalloproteinase inhibitors as a new therapy Nature Medicine 8:35–40. https://doi.org/10.1038/nm0102-35.

7. Ayuso-Sacido A, Moliterno JA, Kratovac S, Kapoor GS, O’Rourke DM, Holland EC, García-Verdugo JM, Roy NS, Boockvar JA (2010) Activated EGFR signaling increases proliferation, survival, and migration and blocks neuronal differentiation in post-natal neural stem cells Journal of Neuro-Oncology 97:323–37. https://doi.org/10.1007/s11060-009-0035-x

8. Baker AT, Zlobin A, Osipo C (2014) Notch-EGFR/HER2 bidirectional crosstalk in breast cancer Frontiers in Oncology 4:360. https://doi.org/10.3389/fonc.2014.00360

9. Bayramli X, Kocagöz Y, Sakizli U, Fuss SH (2017) Patterned arrangements of olfactory receptor gene expression in zebrafish are established by radial movement of specified olfactory sensory neurons Scientific Reports 7:5572. https://doi.org/10.1038/s41598-017-06041-1

10. Beites CL, Kawauchi S, Crocker CE, Calof AL (2005) Identification and molecular regulation of neural stem cells in the olfactory epithelium Experimental Cell Research 306:309–16. https://doi.org/10.1016/j.yexcr.2005.03.027

11. Blobel CP (2005) ADAMs: key components in EGFR signalling and development Nature Review Molecular Cell Biology 6:32–43. https://doi.org/10.1038/nrm1548

12. Buckland ME, Cunningham AM (1998) Alterations in the neurotrophic factors BDNF, GDNF and CNTF in the regenerating olfactory system. Annals of the New York Academy of Sciences 855:260–5. https://doi.org/10.1111/j.1749-6632.1998.tb10579.x

13. Calof AL, Bonnin A, Crocker C, Kawauchi S, Murray RC, Shou J, Wu HH (2002) Progenitor cells of the olfactory receptor neuron lineage Microscopy Research and Technique 58:176–88. https://doi.org/10.1002/jemt.10147

14. Cardone MH, Roy N, Stennicke HR, Salvesen GS, Franke TF, Stanbridge E, Frisch S, Reed JC (1998) Regulation of cell death protease caspase-9 by phosphorylation Science 282:1318–21. https://doi.org/10.1126/science.282.5392.1318

15. Carter LA, MacDonald JL, Roskams AJ (2004) Olfactory horizontal basal cells demonstrate a conserved multipotent progenitor phenotype The Journal of Neuroscience 24:5670–83. https://doi.org/10.1523/JNEUROSCI.0330-04.2004

16. Casas Gimeno G, Paridaen JTML (2022) The symmetry of neural stem cell and progenitor divisions in the vertebrate brain. Frontiers in Cell and Developmental Biology 10:885269. https://doi.org/10.3389/fcell.2022.885269

17. Castanieto A, Johnston MJ, Nystul TG (2014) EGFR signaling promotes self-renewal through the establishment of cell polarity in Drosophila follicle stem cells Elife 3:e04437. https://doi.org/10.7554/eLife.04437

18. Chen X, Fang H, Schwob JE (2004) Multipotency of purified, transplanted globose basal cells in olfactory epithelium The Journal of Comparative Neurology 469:457–74. https://doi.org/10.1002/cne.11031

19. Chen ZH, Luo XC, Yu CR, Huang L (2020) Matrix metalloprotease-mediated cleavage of neural glial-related cell adhesion molecules activates quiescent olfactory stem cells via EGFR Molecular and Cellular Neuroscience 108:103552. https://doi.org/10.1016/j.mcn.2020.103552

20. Clarke MF, Fuller M (2006) Stem cells and cancer: two faces of eve Cell 124:1111–5. https://doi.org/10.1016/j.cell.2006.03.011

21. Clevers H, Loh KM, Nusse R (2014) Stem cell signaling. An integral program for tissue renewal and regeneration: Wnt signaling and stem cell control Science 346:1248012. https://doi.org/10.1126/science.1248012

22. Cunningham F et al. (2022) Ensembl 2022 Nucleic Acids Research, 50:D988–D995. https://doi.org/10.1093/nar/gkab1049

23. Dao DT, Anez-Bustillos L, Adam RM, Puder M, Bielenberg DR (2018) Heparin-binding epidermal growth factor-like growth factor as a critical mediator of tissue repair and regeneration. The American Journal of Pathology 188:2446–2456. https://doi.org/10.1016/j.ajpath.2018.07.016

24. Demirler MC, Sakizli U, Bali B, Kocagöz Y, Eski SE, Ergönen A, Alkiraz AS, Bayramli X, Hassenklöver T, Manzini I, Fuss SH (2020) Purinergic signalling selectively modulates maintenance but not repair neurogenesis in the zebrafish olfactory epithelium The FEBS Journal 287:2699–2722. https://doi.org/110.1111/febs.15170

25. Díaz B, Yuen A, Iizuka S, Higashiyama S, Courtneidge SA (2013) Notch increases the shedding of HB-EGF by ADAM12 to potentiate invadopodia formation in hypoxia Journal of Cell Biology 201:279–92. https://doi.org/10.1083/jcb.201209151

26. Doroquez DB, Rebay I (2006) Signal integration during development: mechanisms of EGFR and Notch pathway function and cross-talk Critical Reviews in Biochemistry and Molecular Biology 41:339–85. https://doi.org/10.1080/10409230600914344

27. Elenius K, Paul S, Allison G, Sun J, Klagsbrun M (1997) Activation of HER4 by heparin-binding EGF-like growth factor stimulates chemotaxis but not proliferation EMBO Journal 16:1268–78. https://doi.org/10.1093/emboj/16.6.1268

28. Falls DL (2003) Neuregulins: functions, forms, and signmaling strategies Experimental Cell Research 284:14–30. https://doi.org/10.1016/s0014-4827(02)00102-7

29. Farbman AI, Buchholz JA (1996) Transforming growth factor-alpha and other growth factors stimulate cell division in olfactory epithelium in vitro Journal of Neurobiology 30:267–80. https://doi.org/10.1002/(SICI)1097-4695(199606)30:2<267::AID-NEU8>3.0.CO;2-3

30. Farbman AI, Ezeh PI (2000) TGF-alpha and olfactory marker protein enhance mitosis in rat olfactory epithelium in vivo Neuroreport 11:3655–8. https://doi.org10.1097/00001756-200011090-00051

31. Féron F, Vincent A, Mackay-Sim A (1999) Dopamine promotes differentiation of olfactory neuron in vitro Brain Research 845:252–9. https://doi.org/10.1016/s0006-8993(99)01959-9

32. Fletcher RB, Prasol MS, Estrada J, Baudhuin A, Vranizan K, Choi YG, Ngai J (2011) p63 regulates olfactory stem cell self-renewal and differentiation Neuron, 72:748–59. https://doi.org10.1016/j.neuron.2011.09.009

33. Fletcher RB, Das D, Gadye L, Street KN, Baudhuin A, Wagner A, Cole MB, Flores Q, Choi YG, Yosef N, Purdom E, Dudoit S, Risso D, Ngai J (2017) Deconstructing olfactory stem cell trajectories at single-cell resolution Cell Stem Cell 20:817–830.e8. https://doi.org/10.1016/j.stem.2017.04.003

34. Fry DW, Kraker AJ, McMichael A, Ambroso LA, Nelson JM, Leopold WR, Connors RW, Bridges AJ (1994) A specific inhibitor of the epidermal growth factor receptor tyrosine kinase Science 265:1093–5. https://doi.org/10.1126/science.8066447

35. Fukuda Y, Katsunuma S, Uranagase A, Nota J, Nibu KI (2018) Effect of intranasal administration of neurotrophic factors on regeneration of chemically degenerated olfactory epithelium in aging mice Neuroreport 29:1400–1404. https://doi.org/10.1097/WNR.0000000000001125

36. Fuller SJ, Sivarajah K, Sugden PH (2008) ErbB receptors, their ligands, and the consequences of their activation and inhibition in the myocardium Journal of Molecular and Cellular Cardiology 44:831–54. https://doi.org/10.1016/j.yjmcc.2008.02.278

37. Gadye L, Das D, Sanchez MA, Street K, Baudhuin A, Wagner A, Cole MB, Choi YG, Yosef N, Purdom E, Dudoit S, Risso D, Ngai J, Fletcher RB (2017) Injury activates transient olfactory stem cell states with diverse lineage capacities. Cell Stem Cell 21:775–790.e9. https://doi.org/10.1016/j.stem.2017.10.014

38. Getchell TV, Narla RK, Little S, Hyde JF, Getchell ML (2000) Horizontal basal cell proliferation in the olfactory epithelium of transforming growth factor-alpha transgenic mice Cell and Tissue Research 299:185–92. https://doi.org/10.1007/s004419900149

39. Gilbert MA, Lin B, Peterson J, Jang W, Schwob JE (2015) Neuregulin1 and ErbB expression in the uninjured and regenerating olfactory mucosa Gene Expression Patterns 19:108–19. https://doi.org/10.1016/j.gep.2015.10.001

40. Gokoffski KK, Wu HH, Beites CL, Kim J, Kim EJ, Matzuk MM, Johnson JE, Lander AD, Calof AL (2011) Activin and GDF11 collaborate in feedback control of neuroepithelial stem cell proliferation and fate Development 138:4131–42. https://doi.org/10.1242/dev.065870

41. Groot AJ, Vooijs MA (2012) The role of Adams in Notch signaling Advances in Experimental Medicine and Biology 727:15–36. https://doi.org/10.1007/978-1-4614-0899-4_2

42. Herrick DB, Lin B, Peterson J, Schnittke N, Schwob JE (2017) Notch1 maintains dormancy of olfactory horizontal basal cells, a reserve neural stem cell Proceedings of the National Academy of Sciences of the United States of America 114:E5589–E5598. https://doi.org/10.1073/pnas.1701333114

43. Hashimoto K, Higashiyama S, Asada H, Hashimura E, Kobayashi T, Sudo K, Nakagawa T, Damm D, Yoshikawa K, Taniguchi N (1994) Heparin-binding epidermal growth factor-like growth factor is an autocrine growth factor for human keratinocytes Journal of Biological Chemistry 269:20060–6. https://doi.org/10.1016/S0021-9258(17)32127-

44. Hasson P, Paroush Z (2006) Crosstalk between the EGFR and other signalling pathways at the level of the global transcriptional corepressor Groucho/TLE British Journal of Cancer 94:771–5. https://doi.org/10.1038/sj.bjc.6603019

45. Herrick DB, Lin B, Peterson J, Schnittke N, Schwob JE (2017) Notch1 maintains dormancy of olfactory horizontal basal cells, a reserve neural stem cell. Proceedings of the National Academy of Sciences of the United States of America 114:E5589–E5598. https://doi.org/10.1073/pnas.1701333114

46. Higashiyama S, Abraham JA, Miller J, Fiddes JC, Klagsbrun M (1991) A heparin-binding growth factor secreted by macrophage-like cells that is related to EGF Science 251:936–9. https://doi.org/10.1126/science.1840698

47. Higashiyama S, Nanba D (2005) ADAM-mediated ectodomain shedding of HB-EGF in receptor cross-talk Biochimica et Biophysica Acta 1751:110–7. https://doi.org/10.1016/j.bbapap.2004.11.009

48. Higashiyama S, Iwabuki H, Morimoto C, Hieda M, Inoue H, Matsushita N (2008) Membrane-anchored growth factors, the epidermal growth factor family: beyond receptor ligands Cancer Science 99:214–20. https://doi.org/10.1111/j.1349-7006.2007.00676.x

49. Holbrook EH, Szumowski KE, Schwob JE (1995) An immunochemical, ultrastructural, and developmental characterization of the horizontal basal cells of rat olfactory epithelium The Journal of Comparative Neurology 363:129–46. https://doi.org/10.1002/cne.903630111

50. Hu T, Li C (2010) Convergence between Wnt-β-catenin and EGFR signaling in cancer Molecular Cancer 9:236. https://doi.org/10.1186/1476-4598-9-236

51. Iqbal T, Byrd-Jacobs C (2010) Rapid degeneration and regeneration of the zebrafish olfactory epithelium after triton X-100 application Chemical Senses 35:351–61. https://doi.org/10.1093/chemse/bjq019

52. Jia C, Doherty JP, Crudgington S, Hegg CC (2009) Activation of purinergic receptors induces proliferation and neuronal differentiation in Swiss Webster mouse olfactory epithelium Neuroscience 163:120–8. https://doi.org/10.1016/j.neuroscience.2009.06.040

53. Jia C, Hegg CC (2012) Neuropeptide Y and extracellular signal-regulated kinase mediate injury-induced neuroregeneration in mouse olfactory epithelium. Molecular and Cellular Neuroscience 49:158–70. https://doi.org/10.1016/j.mcn.2011.11.004

54. Kim EJ, Simpson PJ, Park DJ, Liu BQ, Ronnett GV, Moon C (2005) Leukemia inhibitory factor is a proliferative factor for olfactory sensory neurons Neuroreport 16:25–8. https://doi.org10.1097/00001756-200501190-00007

55. Knight C, James S, Kuntin D, Fox J, Newling K, Hollings S, Pennock R, Genever P (2019) Epidermal growth factor can signal via β-catenin to control proliferation of mesenchymal stem cells independently of canonical Wnt signaling Cellular Signalling 53:256–268. https://doi.org/10.1016/j.cellsig.2018.09.021

56. Kocagöz Y, Demirler MC, Eski SE, Güler K, Dokuzluoglu Z, Fuss SH. (2022) Disparate progenitor cell populations contribute to maintenance and repair neurogenesis in the zebrafish olfactory epithelium Cell and Tissue Research 388:331–358. https://doi.org/10.1007/s00441-022-03597-x

57. Kolev V, Mandinova A, Guinea-Viniegra J, Hu B, Lefort K, Lambertini C, Neel V, Dummer R, Wagner EF, Dotto GP (2008) EGFR signalling as a negative regulator of Notch1 gene transcription and function in proliferating keratinocytes and cancer Nature Cell Biology 10:902–11. https://doi.org/10.1038/ncb1750

58. Korsching SI (2020) Taste and Smell in Zebrafish The Senses: A Comprehensive Reference (Second Edition), Editor(s): Bernd Fritzsch, Elsevier, pp 466–492, ISBN 9780128054093. https://doi.org/10.1016/B978-0-12-809324-5.24155-2

59. Krampera M, Pasini A, Rigo A, Scupoli MT, Tecchio C, Malpeli G, Scarpa A, Dazzi F, Pizzolo G, Vinante F (2005) HB-EGF/HER-1 signaling in bone marrow mesenchymal stem cells: inducing cell expansion and reversibly preventing multilineage differentiation Blood 106:59–66. https://doi.org/10.1182/blood-2004-09-3645

60. Krishna NS, Little SS, Getchell TV (1996) Epidermal growth factor receptor mRNA and protein are expressed in progenitor cells of the olfactory epithelium The Journal of Comparative Neurology 373:297–307. https://doi.org/10.1002/(SICI)1096-9861(19960916)373:2<297::AID-CNE11>3.0.CO;2-I

61. Krolewski RC, Packard A, Schwob JE (2013) Global expression profiling of globose basal cells and neurogenic progression within the olfactory epithelium The Journal of Comparative Neurology 521:833–59. https://doi.org/10.1002/cne.23204

62. Leung CT, Coulombe PA, Reed RR (2007) Contribution of olfactory neural stem cells to tissue maintenance and regeneration Nature Neuroscience 10:720–6. https://doi.org/10.1038/nn1882

63. Li L, Clevers H (2010) Coexistence of quiescent and active adult stem cells in mammals Science 327:542–5. https://doi.org/10.1126/science.1180794

64. Lin B, Coleman JH, Peterson JN, Zunitch MJ, Jang W, Herrick DB, Schwob JE (2017) Injury induces endogenous reprogramming and dedifferentiation of neuronal progenitors to multipotency Cell Stem Cell 21:761–774.e5. https://doi.org/10.1016/j.stem.2017.09.008

65. Linggi B, Carpenter G (2006) ErbB receptors: new insights on mechanisms and biology Trends in Cell Biology 16:649–56. https://doi.org/10.1016/j.tcb.2006.10.008

66. Löffek S, Schilling O, Franzke CW (2011) Biological role of matrix metalloproteinases: a critical balance European Respiratory Journal 38:191–208. https://doi.org/10.1183/09031936.00146510

67. Lucas LM, Dwivedi V, Senfeld JI, Cullum RL, Mill CP, Piazza JT, Bryant IN, Cook LJ, Miller ST, Lott JH 4th, Kelley CM, Knerr EL, Markham JA, Kaufmann DP, Jacobi MA, Shen J, Riese DJ 2nd (2022) The Yin and Yang of ERBB4: Tumor Suppressor and Oncoprotein Pharmacological Reviews 74:18–47. https://doi.org/10.1124/pharmrev.121.000381

68. Lucchesi PA, Sabri A, Belmadani S, Matrougui K (2004) Involvement of metalloproteinases 2/9 in epidermal growth factor receptor transactivation in pressure-induced myogenic tone in mouse mesenteric resistance arteries Circulation 110:3587–93. https://doi.org/10.1161/01.CIR.0000148780.36121.47

69. Mahanthappa NK, Schwarting GA (1993) Peptide growth factor control of olfactory neurogenesis and neuron survival in vitro: roles of EGF and TGF-beta s Neuron 10:293–305. https://doi.org/10.1016/0896-6273(93)90319-m

70. Mitamura T, Higashiyama S, Taniguchi N, Klagsbrun M, Mekada E (1995) Diphtheria toxin binds to the epidermal growth factor (EGF)-like domain of human heparin-binding EGF-like growth factor/diphtheria toxin receptor and inhibits specifically its mitogenic activity Journal of Biological Chemistry 270:1015–9. https://doi.org/10.1074/jbc.270.3.1015

71. Nagata S, Kikuchi M (2020) Emergence of cooperative bistability and robustness of gene regulatory networks PLoS Computational Biology 16:e1007969. https://doi.org/10.1371/journal.pcbi.1007969

72. Nguyen BC, Lefort K, Mandinova A, Antonini D, Devgan V, Della Gatta G, Koster MI, Zhang Z, Wang J, Tommasi di Vignano A, Kitajewski J, Chiorino G, Roop DR, Missero C, Dotto GP (2006) Cross-regulation between Notch and p63 in keratinocyte commitment to differentiation Genes & Development 20:1028–42. https://doi.org/10.1101/gad.1406006

73. Ohta Y, Ichimura K (2000) Immunohistochemical localization of proliferating cells and epidermal growth factor receptors in mouse olfactory epithelium ORL Journal for Oto-Rhino-Laryngology and its Related Specialties 62:20–5. https://doi.org/10.1159/000027710

74. Oka Y, Korsching SI (2011) Shared and unique G alpha proteins in the zebrafish versus mammalian senses of taste and smell Chemical Senses 36:357–65. https://doi.org/10.1093/chemse/bjq138

75. Oka Y, Saraiva LR, Korsching SI (2012) Crypt neurons express a single V1R-related ora gene Chemical Senses 37:219–27. https://doi.org/10.1093/chemse/bjr095

76. Oyagi A, Hara H (2012) Essential roles of heparin-binding epidermal growth factor-like growth factor in the brain. CNS Neuroscience & Therapeutics 18:803–10. https://doi.org/10.1111/j.1755-5949.2012.00371.x

77. Packard A, Schnittke N, Romano RA, Sinha S, Schwob JE (2011) DeltaNp63 regulates stem cell dynamics in the mammalian olfactory epithelium The Journal of Neuroscience 31:8748–59. https://doi.org/10.1523/JNEUROSCI.0681-11.2011

78. Palominos MF, Calfún C, Nardocci G, Candia D, Torres-Paz J, Whitlock KE (2022) The olfactory organ is a unique site for neutrophils in the brain Frontiers in Immunology 13:881702. https://doi.org/10.3389/fimmu.2022.881702

79. Pennock S, Wang Z (2003) Stimulation of cell proliferation by endosomal epidermal growth factor receptor as revealed through two distinct phases of signaling Molecular and Cellular Biology 23:5803–15. https://doi.org/10.1128/MCB.23.16.5803-5815.2003

80. Plendl J, Stierstorfer B, Sinowatz F (1999) Growth factors and their receptors in the olfactory system Anatomia, Histologia, Embryologia 28:73–9. https://doi.org/10.1046/j.1439-0264.1999.00165.x

81. Puschmann TB, Zandén C, Lebkuechner I, Philippot C, de Pablo Y, Liu J, Pekny M (2014) HB-EGF affects astrocyte morphology, proliferation, differentiation, and the expression of intermediate filament proteins Journal of Neurochemistry 128:878–89. https://doi.org/10.1111/jnc.12519

82. R Core Team (2022) R: A language and environment for statistical computing R Foundation for Statistical Computing, Vienna, Austria. https://www.R-project.org

83. Rasmussen HS, McCann PP (1997) Matrix metalloproteinase inhibition as a novel anticancer strategy: a review with special focus on batimastat and marimastat Pharmacology & Therapeutics 75:69–75. https://doi.org/10.1016/s0163-7258(97)00023-5

84. Ruijtenberg S, van den Heuvel S (2016) Coordinating cell proliferation and differentiation: Antagonism between cell cycle regulators and cell type-specific gene expression Cell Cycle 15:196–212. https://doi.org/10.1080/15384101.2015.1120925

85. Sahin U, Weskamp G, Kelly K, Zhou HM, Higashiyama S, Peschon J, Hartmann D, Saftig P, Blobel CP (2004) Distinct roles for ADAM10 and ADAM17 in ectodomain shedding of six EGFR ligands Journal of Cell Biology 164:769–79. https://doi.org/10.1083/jcb.200307137

86. Sato Y, Miyasaka N, Yoshihara Y (2005) Mutually exclusive glomerular innervation by two distinct types of olfactory sensory neurons revealed in transgenic zebrafish The Journal of Neuroscience 25:4889–97. https://doi.org/10.1523/JNEUROSCI.0679-05.2005

87. Schindelin J, Arganda-Carreras I, Frise E, Kaynig V, Longair M, Pietzsch T, Preibisch S, Rueden C, Saalfeld S, Schmid B, Tinevez JY, White DJ, Hartenstein V, Eliceiri K, Tomancak P, Cardona A (2012) Fiji: an open-source platform for biological-image analysis Nature Methods 9:676–82. https://doi.org/10.1038/nmeth.2019

88. Schnittke N, Herrick DB, Lin B, Peterson J, Coleman JH, Packard AI, Jang W, Schwob JE (2015) Transcription factor p63 controls the reserve status but not the stemness of horizontal basal cells in the olfactory epithelium. Proceedings of the National Academy of Sciences of the United States of America 112:E5068–77. https://doi.org/10.1073/pnas.1512272112

89. Schwob JE (2002) Neural regeneration and the peripheral olfactory system The Anatomical Record 269:33–49. https://doi.org/10.1002/ar.10047

90. Schwob JE, Jang W, Holbrook EH, Lin B, Herrick DB, Peterson JN, Hewitt Coleman J (2017) Stem and progenitor cells of the mammalian olfactory epithelium: Taking poietic license The Journal of Comparative Neurology 525:1034–1054. https://doi.org/10.1002/cne.24105

91. She QB, Solit DB, Ye Q, O’Reilly KE, Lobo J, Rosen N (2005) The BAD protein integrates survival signaling by EGFR/MAPK and PI3K/Akt kinase pathways in PTEN-deficient tumor cells Cancer Cell 8:287–97. https://doi.org/10.1016/j.ccr.2005.09.006

92. Simons BD, Clevers H (2011) Strategies for homeostatic stem cell self-renewal in adult tissues Cell 145:851–62. https://doi.org/10.1016/j.cell.2011.05.033

93. Simpson PJ, Wang E, Moon C, Matarazzo V, Cohen DR, Liebl DJ, Ronnett GV (2003) Neurotrophin-3 signaling maintains maturational homeostasis between neuronal populations in the olfactory epithelium Molecular and Cellular Neuroscience 24:858–74. https://doi.org/10.1016/j.mcn.2003.08.001

94. Singh AB, Harris RC (2005) Autocrine, paracrine and juxtacrine signaling by EGFR ligands Cellular Signalling 17:1183–93. https://doi.org/10.1016/j.cellsig.2005.03.026

95. Smedley D, Haider S, Ballester B, Holland R, London D, Thorisson G, Kasprzyk A (2009) BioMart--biological queries made easy BMC Genomics 10:22. https://doi.org/10.1186/1471-2164-10-22.

96. Sundaram MV (2005) The love-hate relationship between Ras and Notch Genes & Development 19:1825–39. https://doi.org/10.1101/gad.1330605

97. Suzuki M, Raab G, Moses MA, Fernandez CA, Klagsbrun M (1997) Matrix metalloproteinase-3 releases active heparin-binding EGF-like growth factor by cleavage at a specific juxtamembrane site Journal of Biological Chemistry 272:31730–7. https://doi.org/10.1074/jbc.272.50.31730

98. Suzuki H, Nikaido M, Hagino-Yamagishi K, Okada N (2015) Distinct functions of two olfactory marker protein genes derived from teleost-specific whole genome duplication BMC Evolutionary Biology 15:245. https://doi.org/10.1186/s12862-015-0530-y

99. Tadeu AM, Horsley V (2013) Notch signaling represses p63 expression in the developing surface ectoderm Development 140:3777–86. https://doi.org/10.1242/dev.093948

100. Umata T, Hirata M, Takahashi T, Ryu F, Shida S, Takahashi Y, Tsuneoka M, Miura Y, Masuda M, Horiguchi Y, Mekada E (2001) A dual signaling cascade that regulates the ectodomain shedding of heparin-binding epidermal growth factor-like growth factor Journal of Biological Chemistry 276:30475–82. https://doi.org/10.1074/jbc.M103673200

101. Visvader JE, Clevers H (2016) Tissue-specific designs of stem cell hierarchies Nature Cell Biology 18:349–55. https://doi.org/10.1038/ncb3332

102. Wakisaka N, Miyasaka N, Koide T, Masuda M, Hiraki-Kajiyama T, Yoshihara Y (2017) An adenosine receptor for olfaction in fish Current Biology 27:1437–1447.e4. https://doi.org/10.1016/j.cub.2017.04.014

103. Wan J, Ramachandran R, Goldman D (2012) HB-EGF is necessary and sufficient for Müller glia dedifferentiation and retina regeneration Developmental Cell 22:334–47. https://doi.org/10.1016/j.devcel.2011.11.020

104. Wan J, Zhao XF, Vojtek A, Goldman D (2014) Retinal injury, growth factors, and cytokines converge on β-catenin and pStat3 signaling to stimulate retina regeneration Cell Reports 9:285–297. https://doi.org/10.1016/j.celrep.2014.08.048

105. Wang YZ, Yamagami T, Gan Q, Wang Y, Zhao T, Hamad S, Lott P, Schnittke N, Schwob JE, Zhou CJ (2011) Canonical Wnt signaling promotes the proliferation and neurogenesis of peripheral olfactory stem cells during postnatal development and adult regeneration Journal of Cell Science 124:1553–63. https://doi.org/10.1242/jcs.080580

106. Wang YX, Feige P, Brun CE, Hekmatnejad B, Dumont NA, Renaud JM, Faulkes S, Guindon DE, Rudnicki MA (2019) EGFR-Aurka signaling rescues polarity and regeneration defects in dystrophin-deficient muscle stem cells by increasing asymmetric divisions Cell Stem Cell 24:419–432.e6. https://doi.org/10.1016/j.stem.2019.01.002

107. Wee P, Wang Z (2017) Epidermal growth factor receptor cell proliferation signaling pathways Cancers 9:52. https://doi.org/10.3390/cancers9050052

108. Wojtowicz-Praga SM, Dickson RB, Hawkins MJ (1997) Matrix metalloproteinase inhibitors Investigational New Drugs 15:61–75. https://doi.org/10.1023/a:1005722729132

109. Wu HH, Ivkovic S, Murray RC, Jaramillo S, Lyons KM, Johnson JE, Calof AL (2003) Autoregulation of neurogenesis by GDF11 Neuron 37:197–207. https://doi.org/10.1016/s0896-6273(02)01172-8

110. Yan Y, Shirakabe K, Werb Z (2002) The metalloprotease Kuzbanian (ADAM10) mediates the transactivation of EGF receptor by G protein-coupled receptors Journal of Cell Biology 158:221–6. https://doi.org/10.1083/jcb.200112026

111. Zhang Y, Sui F, Ma J, Ren X, Guan H, Yang Q, Shi J, Ji M, Shi B, Sun Y, Hou P (2017) Positive feedback loops between NrCAM and major signaling pathways contribute to thyroid tumorigenesis The Journal of Clinical Endocrinology and Metabolism 102:613–624. https://doi.org/10.1210/jc.2016-1677

112. Zhao XF, Wan J, Powell C, Ramachandran R, Myers MG Jr, Goldman D (2014) Leptin and IL-6 family cytokines synergize to stimulate Müller glia reprogramming and retina regeneration Cell Reports 9:272–284. https://doi.org/10.1016/j.celrep.2014.08.047

113. Zhou L, Wang DS, Li QJ, Sun W, Zhang Y, Dou KF (2012) Downregulation of the Notch signaling pathway inhibits hepatocellular carcinoma cell invasion by inactivation of matrix metalloproteinase-2 and -9 and vascular endothelial growth factor Oncology Reports 28:874–82. https://doi.org/10.3892/or.2012.1880

